# Tracking *Pseudomonas aeruginosa* transmissions due to environmental contamination after discharge in ICUs using mathematical models

**DOI:** 10.1101/492538

**Authors:** Thi Mui Pham, Mirjam Kretzschmar, Xavier Bertrand, Martin Bootsma, on behalf of COMBACTE-MAGNET Consortium

**Affiliations:** Julius Center for Health Sciences and Primary Care of the UMC Utrecht, Utrecht, The Netherlands; Mathematical Institute, Utrecht University, Utrecht, The Netherlands; Centre Hospitalier Universitaire Besançon, Besançon, France; National Institute for Public Health and the Environment (RIVM), Utrecht, The Netherlands

## Abstract

*Pseudomonas aeruginosa* (*P. aeruginosa*) is an important cause of healthcare-associated infections, particularly in immunocompromised patients. Understanding how this multi-drug resistant pathogen is transmitted within intensive care units (ICUs) is crucial for devising and evaluating successful control strategies. While it is known that moist environments serve as natural reservoirs for *P. aeruginosa*, there is little quantitative evidence regarding the contribution of environmental contamination to its transmission within ICUs. Previous studies on other nosocomial pathogens rely on deploying specific values for environmental parameters derived from costly and laborious genotyping. Using solely longitudinal surveillance data, we estimated the relative importance of *P. aeruginosa* transmission routes by exploiting the fact that different routes cause different pattern of fluctuations in the prevalence.

We developed a mathematical model including endogenous colonization, cross-transmission and environmental contamination. Patients contribute to a pool of pathogens by shedding bacteria to the environment. Natural decay and cleaning of the environment lead to a reduction of that pool. By assigning the bacterial load shed during an ICU stay to cross-transmission, we were able to disentangle environmental contamination during and after a patient’s stay. Based on a data-augmented Markov Chain Monte Carlo method the relative importance of the considered acquisition routes is determined for two ICUs of the University hospital in Besancon (France). We used information about the admission and discharge days, screening days and screening results of the ICU patients.

Both cross-transmission and endogenous transmission play a significant role in the transmission process in both ICUs. In contrast, only about 1% of the total transmissions were due to environmental contamination after discharge.

Improved cleaning of the environment after discharge would have only a limited impact regarding the prevention of *P.A*. infections in the two considered ICUs. Our model was developed for *P. aeruginosa* but can be easily applied to other pathogens as well.

**Author summary:** Understanding the transmission dynamics of multi-drug resistant pathogens in intensive-care units is essential for designing successful infection control strategies. We developed a method that estimates the relative importance of several transmission routes of *Pseudomonas aeruginosa* (*P. aeruginosa*), a bacterium intrinsically resistant to multiple antibiotics and known to be a major contributor to hospital-acquired infections. Our model includes three different routes: endogenous colonization, cross-transmission and environmental contamination. Since moist environments may serve as natural reservoirs for *P. aeruginosa*, we focused our study on environmental contamination. Patients contribute to a pool of pathogens by shedding bacteria to the environment. Natural decay and cleaning of the environment lead to a reduction of that pool. By assigning the bacterial load shed during an ICU stay to cross-transmission, we disentangled environmental contamination during and after a patient’s stay. Previous studies were able to assess the role of environmental contamination for specific hospitals using laborious and costly genotyping methods. In contrast, we only used surveillance screening data to estimate the relative contributions of the considered transmission routes. Our results can be used to tailor or assess the effect of interventions. Our model was developed for *P. aeruginosa* but can be easily applied to other pathogens as well.

## Introduction

Hospital-acquired infections are a major cause of morbidity and mortality worldwide [1]. In industrialized countries, about 5 – 10% of admitted acute-care patients are affected whereas the risk is even higher in developing countries [2].

Due to its intrinsic resistance to multiple antibiotics, *Pseudomonas aeruginosa* (short *P. aeruginosa* or *P. A.*) is an important contributor to nosocomial infections [3–5]. The most serious *P. aeruginosa* infections lead to bacteremia, pneumonia, urosepsis, wound infection as well as secondary infection of burns [6]. In 2018, the World Health Organization has recognized *P. aeruginosa* as a serious health-care threat by including it in the list of antibiotic-resistant highest priority pathogens [7].

Given the severe consequences of *P. aeruginosa* infections, in particular for critically-ill patients, it is clear that strategies preventing infections are seen as a key priority. However, infections are recognized as only the tip of the iceberg, while colonizations represent the true load of pathogens carried by patients in the intensive-care unit (ICU). Understanding the dynamics of *P. aeruginosa* colonizations is therefore crucial for developing and evaluating infection control policies.

There are several modes of transmission for colonizations. The endogenous route, due to e. g. antibiotic selection pressure, was regarded as the most important route of *P. aeruginosa* [8,9], but, more and more evidence has emerged on the importance of exogenous sources. Cross-transmission usually caused by temporarily contaminated hands of health-care workers (HCWs) has been identified as an additional source of transmission [10–12]. It is furthermore known that moist environments (e.g. soil and water) may serve as natural reservoirs of *P. aeruginosa* and that it can persist for months on dry inanimate surfaces [13]. Thus, an important question emerged: Does the contamination of the environment with *P. aeruginosa* strains have a substantial impact on the transmission within ICUs even after the discharge of patients?

Quantifying the relative importance of routes of transmission may serve as an essential tool in answering this question. There is little quantitative evidence in the scientific literature regarding the importance of environmental contamination within the transmission dynamics of *P. aeruginosa* especially for non-epidemic situations. Prior studies analyzing the importance of contaminated surfaces on the transmission of other nosocomial pathogens have been conducted, e.g., for Methicillin-resistant Staphylococcus aureus (MRSA) and Vancomycin-Resistant Enterococci (VRE) [14–18]. However, they rely on deploying specific values for model parameters corresponding to the environment. Such information was obtained from previous studies that conducted extensive epidemiological surveillance in combination with costly, laborious as well as time-consuming methods of genotyping. Thus, these methods cannot be easily applied to other nosocomial pathogens without this cumbersome preliminary work.

In this paper, we present a mathematical transmission model that includes, besides endogenous and cross-transmission, environmental contamination as an additional route. Our primary aim is to estimate the relative contributions of the three transmission routes using only longitudinal prevalence data as input. In particular, we are interested in the relative importance of environmental contamination after discharge. We used data from two ICUs of the University hospital in Besancon to estimate the parameters that characterize the transmission routes. The estimation procedure is based on a data-augmented Markov chain Monte Carlo simulation [19]. To our knowledge, this is the first quantitative analysis of the impact of environmental contamination after discharge on *P. aeruginosa* transmissions in ICUs using solely routine surveillance data.

## Materials and methods

In this section, we present our framework for modeling the transmission routes of *P. aeruginosa* including environmental contamination, as well as the method for computing the relative contributions of the routes. We further elaborate on the procedure that we used to estimate the relevant transmission parameters. A brief introduction to the data used for the analysis is given. We describe the model selection as well as model assessment procedures that are used to compare the developed models and to assess the model fit to the data.

### Transmission models

The underlying model for our algorithm is a compartmental Si-model (e.g. [20]). All patients are admitted to an ICU and either belong to the susceptible (*P. aeruginosa* negative) or colonized (*P. aeruginosa* positive) compartment at any given time. The latter includes patients with asymptotic carriage and those with *P. aeruginosa* infection. A susceptible patient may become colonized at a certain transmission rate, which depends on the colonization pressure in the ward at the time.

A stochastic model was developed to reflect the transmission process. The model incorporates three different modes of transmission through which colonization can be acquired. They are distinguished based on the different patterns in the prevalence time series induced by each of them. The endogenous route is independent of other patients and is represented as a constant rate. Consequently, the corresponding prevalence is expected to fluctuate around the mean value and the probability of acquisition for an uncolonized patient is constant during the time period. Sources may be the introduction through visitors or permanently contaminated environments, such as sinks or air-conditioning. Cross-transmission, usually occurring via temporarily contaminated hands of health-care workers, is proportional to the fraction of colonized patients in the wards. The probability of colonization due to cross-transmission is high if the number of colonized patients is high and vice versa. Environmental contamination is modeled on a ward-level represented as a general pool of bacteria linked to objects contaminated by colonized patients. Bacterial load may persist in the environment even after the discharge of patients. This leads to higher probabilities of acquiring colonization after outbreaks, even when the number of colonized patients is low.

The force of infection *λ*(*t*), i.e. the probability per unit of time *t* for a susceptible patient to become colonized, is modeled as

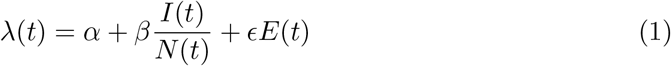

where *I*(*t*) is the number of colonized patients, *N*(*t*) the total number of patients and *E*(*t*) is a compartment tracking the bacterial load present in the ward at time *t*. The parameters *α*, *β* and *ε* are transmission parameters linked to the endogenous term, fraction of colonized patients and the environmental bacterial load, respectively. A schematic of the transmission model is presented in Fig 1.

**Fig 1.**
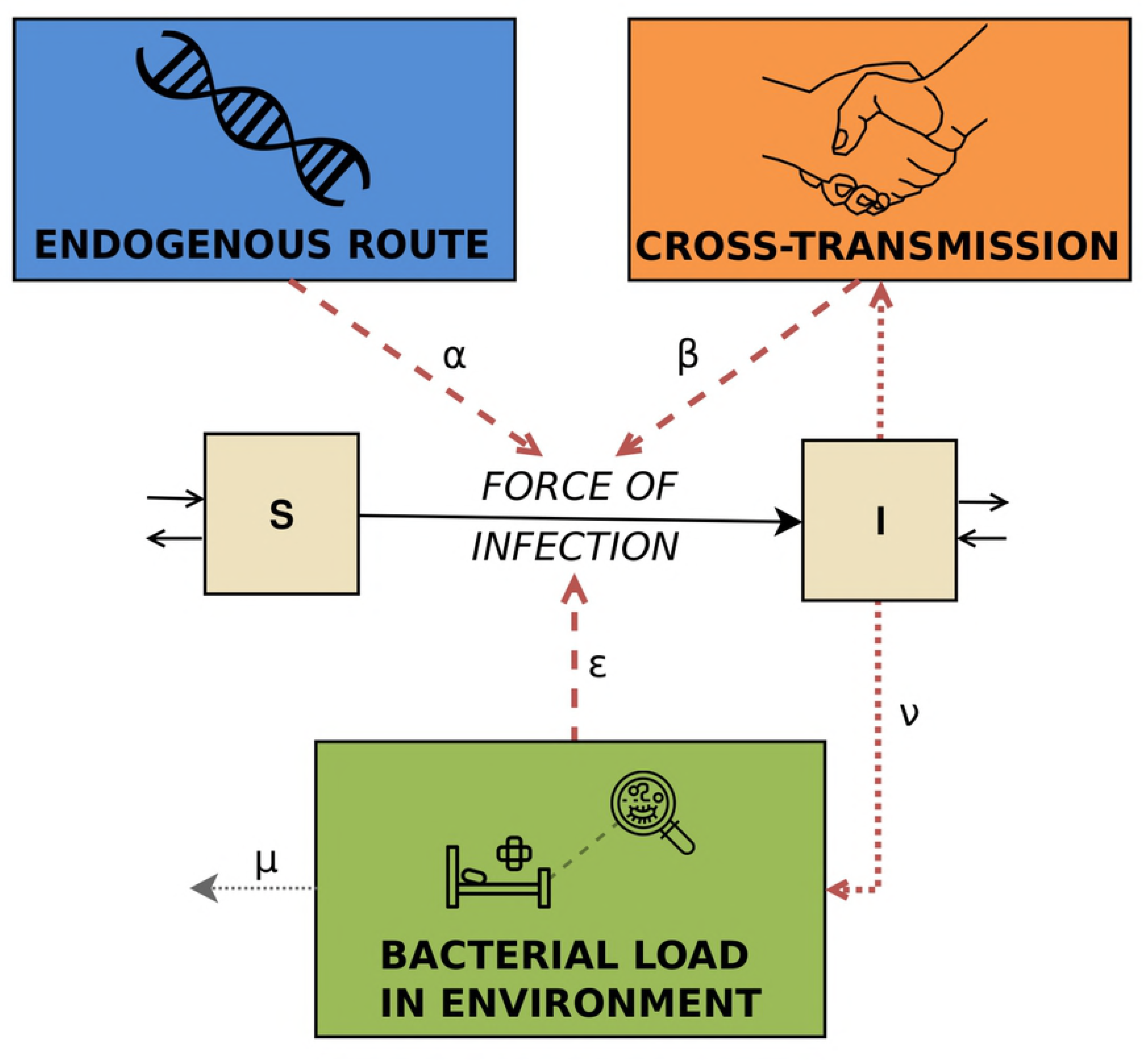
Schematic of the full transmission model. It represents the three different routes, i.e. endogenous route, cross-transmission and environmental contamination.

The described model is subject to the following further assumptions:

- Once colonized, patients remain colonized during the rest of the stay. This assumption is appropriate when the average length of stay of patients does not exceed the duration of colonization, as is the case for *P. aeruginosa*.
- Colonization is assumed to be undetectable until a certain detectable bacterial level is reached. We do not distinguish between several levels of colonization. Furthermore, the detection of carriage in specimen is assumed to be the same for each screening separately.
- Assuming that HCWs are colonized for a short period of time (typically until the next disinfection) in comparison with the length of carriage for patients, we use a quasi-steady state approximation [20]. This means that contact patterns between patients and HCWs are not explicitly modeled and we assume direct patient-to-patient transmission.
- All strains of *P. aeruginosa* are assumed to have the same transmission characteristics. We therefore assume that all colonized patients may be a source of transmission and contribute equally to the colonization pressure.
- All susceptible patients are assumed to be equally susceptible.

In order to analyze the impact of environmental contamination after the discharge of colonized patients, we model the underlying mechanism leading to the presence of pathogens in the environment after discharge. Patients contribute to the bacterial load by shedding *P. aeruginosa* at a rate *ν* during their stay. Furthermore, natural clearance and cleaning lead to a reduction of *P. aeruginosa* bacteria in the environment at a rate *μ*. The change of environmental contamination can be described by

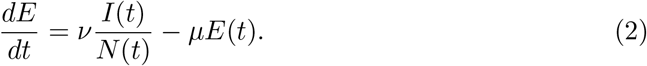

The differential equation (2) is solved by assuming *I*(*t*) = *I_t_* and *N*(*t*) = *N_t_* are known piece-wise constant functions with steps at times *t*_0_, *t*_1_, …, *t*_*N*_. For *t*_*i*_ ∈ {*t*_0_,…, *t*_*N*_}, it holds

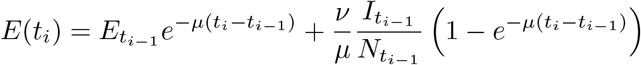

while

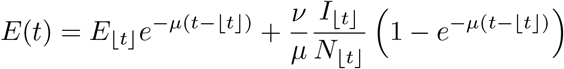

for ⎿*t*⏌:= max{*x* ∈ {*t*_0_, …, *t*_*N*_} | *x* ≤ *t*} and *t* ∈ ℝ \ {*t*_0_, *t*_1_, …, *t*_*N*_} (full details are given in S1 Text). The initial amount of bacterial load is denoted by *E*_0_:= *E*(*t*_0_). Given the number of colonized patients at a certain time *t*, the bacterial load *E*(*t*) is deterministic. The acquisitions are stochastic based on the force of infection in (1). Under the assumption of a force of infection *λ*(*x*) at time *x*, the cumulative probability of any given susceptible person of becoming colonized in [0, *t*] is 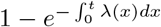 (see e.g. [21]).

All parameters, namely *α*, *β*, *ε*, *μ*, *ν* and *E*_0_ are assumed to be non-negative. By setting certain transmission parameters (*α*, *β* or *ε*) to zero, model variants may be defined. In this paper, we additionally consider a submodel with *ε* = 0, where environmental contamination is not explicitly modeled and therefore only two transmission routes are considered. The force of infection for this transmission model with two acquisition routes is then given by 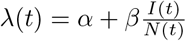.

### Relative contributions of transmission routes

For the prevention of colonization or infection with *P. aeruginosa*, specific intervention control strategies can be designed dependent on the relative importance of the transmission routes. However, for each observed acquisition of colonization, the responsible transmission route is unknown. And yet, for every acquisition, the probability that the colonization was due to a certain route can be determined given that parameter values, the level of environmental contamination and the number of colonized patients are known. Thus, by estimating the transmission parameters *α*, *β*, *ε*, *μ* and *ν*, we were able to approximate the relative contributions of each transmission route to the total number of acquisitions.

The probability of acquisition can be approximated by the force of infection. It consists of different terms that can be assigned to the transmission routes under consideration, i.e.

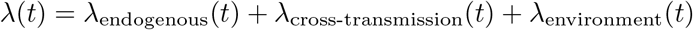

The primary aim of this paper is to estimate the relative contribution of environmental contamination after discharge in order to estimate the role of terminal environmental cleaning among ICU patients. According to our full model, bacterial load is produced by a colonized patient currently present. The cumulative bacterial load increases over time until the respective patient is discharged. After discharge, shedding of that particular patient stops and decreases over time. The bacterial load shed during a patient’s stay (which may then be transmitted via HCWs to other patients) is assigned to cross-transmission as in practice, it may not be distinguished from the classical definition of cross-transmission. The bacterial load persisting after discharge is the variable of interest and represents the impact of an already discharged patient on the current transmissions in the ICU. A schematic of the bacterial load of a single patient over time and its attribution to the different transmission routes is visualized in Fig 2.

**Fig 2.**
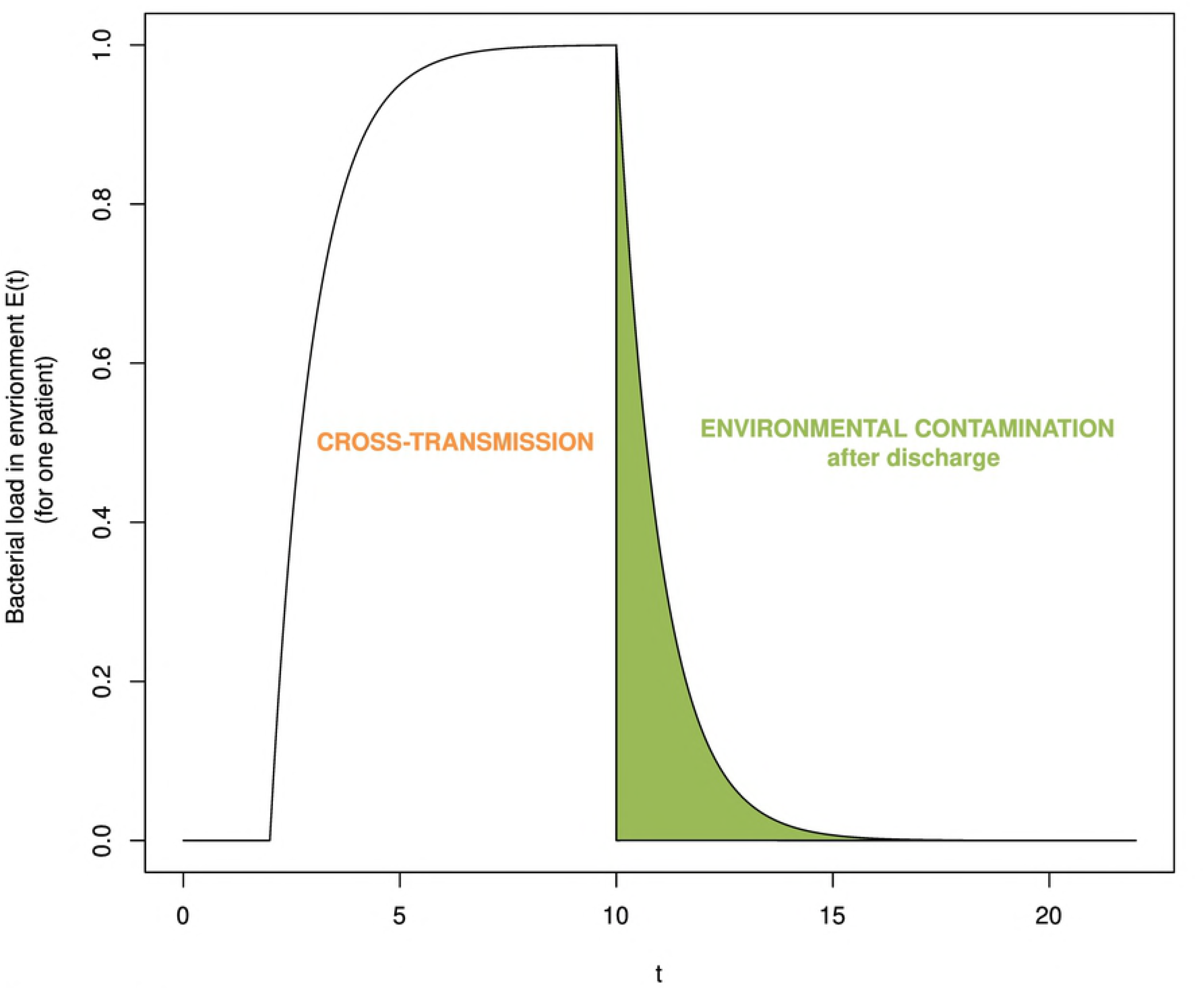
Schematic of the bacterial load shed by a patient developing over time. The bacterial load that is shed during a patient’s stay is assigned to cross-transmission. Environmental contamination after discharge accounts only for the bacterial load persisting after the discharge of that patient.

The previous explanation leads to the following attribution of the terms to the different acquisition routes

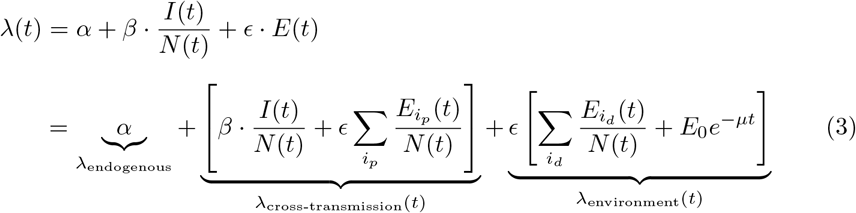

where *i*_*p*_ indicates a colonized patient that is present at time *t* and *i*_*d*_ a colonized patient that has been colonized prior to *t* but was already discharged. The bacterial load produced by patient *i* at time *t* is given by

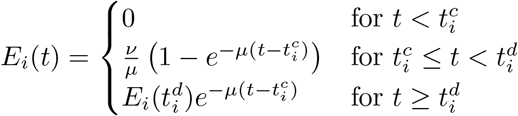

where 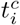 is the time of colonization and 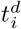 the time of discharge of patient *i*.

In continuous time, the relative contribution of a specific route to the overall number of acquired colonizations is determined by the ratio of the probability of colonization due to that route and the probability of colonization:

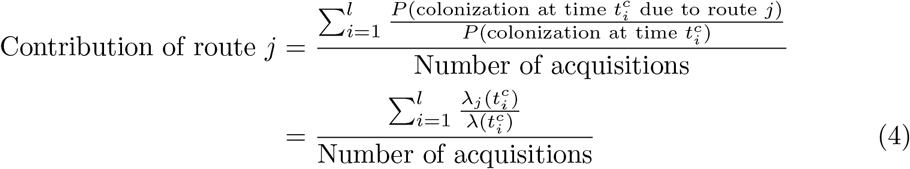

where *l* is the number of colonized patients, 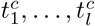 represent the times of colonization and *j* can be either of the three considered routes. The relative contributions are then given by:

- Contribution of endogenous route = 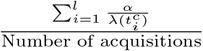
- Contribution of cross-transmission = 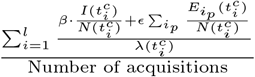
- Contribution of environmental contamination = 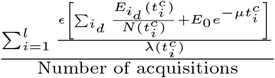

For the submodel including only endogenous and cross-transmission, the computation of the relative contribution is derived from above by setting *ε* = 0.

More details on the calculations can be found in S3 Text. In practice, colonization events are observed only in discrete times. The formulas for the transmission model and the relative contribution are adapted for this discrete time assumption and are elaborated in S2-S3 Texts. Since the calculations for the relative contributions of the transmission routes in the discrete-time scenario require the use of the gamma function and therefore become computationally intensive, we use the continuous-time formulas as approximations. Since values of the force of infection *λ*(*t*) are typically small (< 0.25), the force of infection itself is a good approximation of the probability of infection as 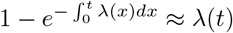 for small values of *λ*(*t*). Hence, the discrete-time formulas for the relative contributions can be approximated by the continuous-time formulas evaluated at discrete time steps.

### Estimation procedure

We assume that a patient is admitted to the ICU at time 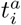 and discharged at time 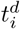. The probability that a patient is admitted already colonized is described by the importation probability *f*. The rate at which a susceptible patient transitions to being colonized is given by Eq. (1). The colonization state of an individual patient is determined from screening information. We suppose that for each patient *i* a set of screening results 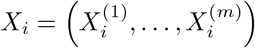, taken on days 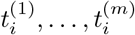 is available. The set of all screening results is denoted by *X* = {*X*_1_,…, *X_n_*} where *n* is the total number of patients. Since screening tests are typically intermittent and imperfect, we define the test sensitivity *ϕ*, i.e. probability that a colonized patient has a positive result.

The aim is to estimate the model parameters *α*, *β*, *ε*, *μ, ν* and *E*_0_ as well as the sensitivity of the screening test *ϕ* and the importation rate *f* based on longitudinal data. The relative contributions of the transmission routes can then be estimated following the description in (4). The key idea of the estimation procedure is to fit a stochastic transmission model to the observed data. It is based on certain patterns of fluctuations in the prevalence linked to the different transmission routes (as previously described in section *Transmission models*).

In the analysis, we use the following input data for each patient:

- day of admission
- day of discharge
- screening days and results.

Thus, we use a day as the smallest time unit in our model and assume that events occur in daily intervals. In principle, other time units may be chosen for an analysis if the required information on admission, discharge and culturing is available. However, smaller units may increase the computational time.

If transmission dynamics were perfectly observed, it would be straightforward to calculate the likelihood of the data given parameters *θ* = {*α*, *β*, *ε*, *μ*, *ν*, *ϕ*, *f*}. However, the true colonization time of a patient is typically unobserved which leads to uncertainty about the true prevalence at any given time. Hence, the likelihood is analytically intractable. We adapted the data-augmented Markov-chain Monte Carlo (MCMC) algorithm (based on [19]) to estimate the posterior distributions of the model parameters. At each iteration, imperfectly observed colonization times are imputed and model parameters *θ* sampled. This method accounts for imperfect and infrequent screening, missing admission and discharge swabs and leads to an estimation of the true (rather than the observed) prevalence on admission. Precise details of the analysis can be found in S5 Text. The algorithm was implemented in C++ and was tested using simulated data. Convergence of the MCMC chains were verified using visual inspection. We used uninformative exponential priors Exp(0.001) for the transmission parameters *α*, *β*, *ε* and *μ*. Parameters for the proposal distribution were tuned in order to ensure rapid convergence. Similar to [26], we estimated the sensitivity *ϕ* and importation parameter *f* using uninformative beta prior distributions Beta(1,1). The initial bacterial load *E*_0_ was approximated by 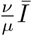 with 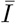 being the mean prevalence in the ward.

The MCMC algorithm was run for 500,000 iterations following a burn-in of 30,000 iterations. The MCMC iterations were then thinned by a factor of 10, leaving 50, 000 iterations for inference. In each iteration, 20 data-augmentation steps were performed with each augmentation chosen at random.

During the estimation process, several assumptions are made.

- Incorporating both sensitivity and specificity parameters in a model may cause identifiability issues. Thus, test specificity was assumed to be 100%, meaning that positive results were assumed to be true positive. Experimental results indicate the specificity of screening tests to be close to 100% [22].
- The initial bacterial load *E*_0_ is assumed to be the environmental contamination at the beginning of the study period. The effect of *E*_0_ diminishes proportionally to exp(–*μ*) per day. It is therefore sufficient to use an approximation rather than including it as a parameter in the estimation process. We use the equilibrium state of (2) as an approximation, i.e.

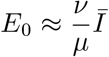

where 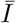 represents the mean prevalence in the ward. The environmental contribution to the force of infection at time *t* is *ε* · *E*(*t*). As the total amount of environmental contamination *E*(*t*) is unobserved, it is only possible to estimate the product *ε* · *E*(*t*) and therefore, we assume the shedding parameter *ν* to be fixed at 0.1. All remaining parameters are to be estimated from the data.
- Colonization was defined as the presence of bacteria at the screening sites as reported in the available data. Admission and screening are assumed to occur at 12:00pm and discharge at 11:59am.
- Re-admissions are not accounted for. Instead every new admission is treated as a new patient. The probability to be positive on admission is therefore identical for all patients, irrespective whether it is a readmission or not. Since we are interested in the overall prevalence and overall relative contribution of the acquisition routes rather than individual predictions, we do not expect this to have a major influence on our results.
- Since the smallest time unit is one day, colonization events occurring on a particular day are assumed to be independent.
- A negative result on the day of colonization is considered to be a false negative result.
- It is assumed that colonized patients contributed to the total colonized population from the day after colonization onwards, or for importations, from the day of admission. This assumption leads to an underestimation of the number of acquisitions for colonization times at the beginning of the day (but just after screening). On the other hand, since pathogenic bacteria such as P. aeruginosa undergo a *lag phase* during their growth cycle, in which the bacteria adapt to the new environment and are not yet able to divide, onward transmission events are likely to be rare during the early stages of colonization. Therefore, the number of onward transmissions are likely to be overestimated for colonizations occurring at the end of the day.

### Data

The data used in the current analysis were collected from two ICUs, denoted by A and B, between 1999 and 2017 at University Hospital of Besancon, eastern France, in the framework of a systematic screening for *P.aeruginosa*. The data sets include admission and discharge dates as well as dates, sites and results of culturing of adult patients. ICU A is a surgical ICU that comprised 15 beds in the time period 1999-2008 and 20 from 2010 till 2017. The ICU was renovated between 2008 and 2009 and the number of beds was increased after completion of the renovation work. ICU B, a non-surgical ICU, had 15 beds from 2000 till 2011 and increased to 20 beds afterwards. Rectal and nose swabs were obtained upon admission (during the first 48 hours) and once a week thereafter. A positive result on one of the swabs was counted as a positive culture. A negative culture resulted from a negative culture on both swabs taken at the specific day. More than 84% of admitted patients were screened. As HCWs, including physicians, were (with minor exceptions) working only in one of the ICUs during the whole study period, the two ICUs can be treated independently in the analysis.

Since 2000, the hand hygiene procedures recommended in both ICUs is rubbing with alcohol-based gels, or solutions (ABS). Cleaning of the rooms is done daily by using the detergent-disinfectant Aniosurf^®^. The sinks were cleaned daily before pouring the detergent-disinfectant Aniosurf^®^ into the U-bends. Plumbing fittings were descaled weekly.

In our main analysis, data for each ICU and each time period (before and after renovation) was treated as distinctive data sets, resulting in four different analyses. No pooling of the results were performed. In a second analysis, the data for the different time periods and different ICUs were combined. The results are compared with the main analysis and are presented in S9-S10 Tables. Each data set was analyzed using

- the full model including endogenous, cross-transmission and environmental contamination after discharge,
- the submodel with only endogenous and exogenous transmission.

Patient data were anonymized and de-identified prior to analysis.

### Model selection

To assess the relative performance of a given model, we used a version of the deviance information criterion (DIC) based on [23]. For an estimated parameter set *θ* and observed data set *x* it is computed as the expected deviance plus the effective number of parameters: 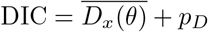. A lower value indicates a better fit. The effective number of parameters *p*_*D*_ represents a complexity measure and is calculated by the difference of the posterior mean deviance and the deviance at the posterior mean: 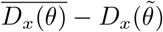. In this paper, we use the approximation 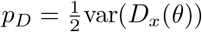 introduced by [23].

The DIC is a simple measure that can be used to compare hierarchical models. Furthermore, it allows determining whether two data sets may be concatenated or should be treated separate. The idea is to distinguish two models: one that includes one parameter set for both ICUs (and therefore treats them as concatenated) and one that includes different parameter sets for each ICU (and thus treats them as separate). The first scenario leads to one analysis and one DIC value whereas the second model results in two independent analyses and hence two DIC values. The sum of the DICs of the latter may be compared to the DIC value of the first scenario. A smaller DIC value is preferred. More details can be found in S6 Text.

### Model assessment

We chose to check the adequacy of the models using the following approach. The ability of the model to predict the probability of acquisition based on the predicted force of infection was assessed. The computed numerical values for the force of infection are assigned to a bin representing the segment covering the numerical value. For a given value *λ* of the force of infection, the theoretical probability of acquisition *p*_acq_ per susceptible patient is computed by 1 – exp(–*λ*). The predicted fraction of acquisitions *f*_acq_ is computed by dividing the number of acquisitions *N*_acq_ by the number of susceptible patients *N*_susc_. We compute 95% confidence intervals assuming that the number of acquisitions follows a binomial distribution of Bin(*N*_susc_, *f*_acq_). The described method is performed for 100 MCMC updates. Coverage probabilities are computed to determine the actual proportion of updates for which the interval contains the theoretical probability of acquisition. We set the nominal confidence level to 0.95. A good fit is given when the actual coverage probability is (more or less) equal to the nominal confidence level. In order to avoid the coverage probability tending to zero when *p*_acq_ tends to 0 or 1, Jeffreys confidence intervals are used (as recommended in [24]). When *N*_acq_ = 0 the lower limit is set to 0, and when *N*_acq_ = *N*_susc_ the upper limit is set to 1.

## Results

### Descriptive analysis of data

The descriptive statistics of the data sets corresponding to ICU A and B with respect to the number of admissions, lengths of stay and colonization characteristics are shown in Table 1. The time period referred to as *before renovation* (short before) is defined as 20/04/1999 – 20/05/2008 (approx. 9 years) for ICU A and 11/01/2000 – 12/01/2011 (approx. 11 years) for ICU B. The time period referred to as *after renovation* (short after) is defined as 21/05/2008 – 21/03/2017 (approx. 7.2 years) for ICU A and 13/01/2011 – 13/09/2016 (approx. 5.8 years) for ICU B. In ICU A, the number of beds decreased during the renovation. Hence, we decided to remove the renovation period from the analysis for ICU A.

**Table 1.** Descriptive statistics for the *P. aeruginosa* carriage data collected from two ICUs at University Hospital Besangon, France, 1999-2017.

|  | No. |  | Median (IQR)<sup>†</sup> |  | % |  |
| --- | --- | --- | --- | --- | --- | --- |
|  | BEFORE<sup>‡</sup> | AFTER<sup>§</sup> | BEFORE | AFTER | BEFORE | AFTER |
| **ICU A** |  |  |  |  |  |  |
| Study length, days | 3200 | 2340 |  |  |  |  |
| Readmissions | 367 | 463 |  |  | 9.3 | 11.8 |
| Admissions* | 3261 | 3398 |  |  |  |  |
| Length of stay, days |  |  | 8.0 (3.0-19.0) | 8.0 (3.0-16.0) |  |  |
| Patients positive on admission<sup>¶</sup> | 69 | 104 |  |  | 3.5 | 3.7 |
| Observed P.A. acquisitions<sup>\|\|¶</sup> | 363 | 289 |  |  | 11.1 | 8.5 |
| **ICU B** |  |  |  |  |  |  |
| Study length, days | 4020 | 2079 |  |  |  |  |
| Readmissions | 891 | 1968 |  |  | 15.4 | 17.8 |
| Admissions | 4360 | 3384 |  |  |  |  |
| Length of stay, days |  |  | 8.0 (3.0-18.0) | 8.0 (3.0-15.0) |  |  |
| Patients positive on admission | 168 | 175 |  |  | 4.7 | 6.3 |
| Observed P.A. acquisitions | 569 | 423 |  |  | 13.1 | 12.5 |

\*Each ICU stay was counted separately, even in case of a multiple ICU stay within a given hospitalization.
<sup>†</sup> Interquartile range
<sup>‡</sup> 20/04/1999 – 23/01/2008 for ICU A and 11/01/2000 – 12/01/2011 ICU B.
<sup>§</sup> 06/11/2010 – 21/03/2017 for ICU A and 13/01/2011 – 13/09/2016 for ICU B.
<sup>¶</sup> False negative results are not taken into account.
<sup>||</sup> An acquisition is when a patient test negative on admission and had a postive result before discharge.

In total, 10,378 patients (5891 admitted to ICU A and 4487 to ICU B) and 37,638 screening results (14,631 in ICU A and 23,007 in ICU B) were included in the analysis. The number of readmissions is higher for ICU B than for ICU A. In our analysis, every admission was treated separately (as a new patient) resulting in 14,403 admissions (6,659 admitted to ICU A and 7,744 to ICU B).

The corresponding median length of stay was 8.0 days for both ICUs before and after renovation, respectively. Hence, there is hardly any difference between the ICUs, nor between the two time periods regarding the median length of stay.

In both ICUs, the fraction of patients who were positive on admission was slightly higher after renovation. In contrast, the observed fraction of patients who acquired colonization slightly decreased after renovation. There were 1,644 patients (652 in ICU A and 992 in ICU B) observed to be colonized during their stay and 516 patients (173 in ICU A and 343 in ICU B) observed to be colonized on admission. The percentage of patients admitted positively on admission and with acquired colonization is higher in ICU B than in ICU A. The total number of patients per ICU and the number of positive cultures are visualized in Figs 3 and 4.

**Fig 3.**
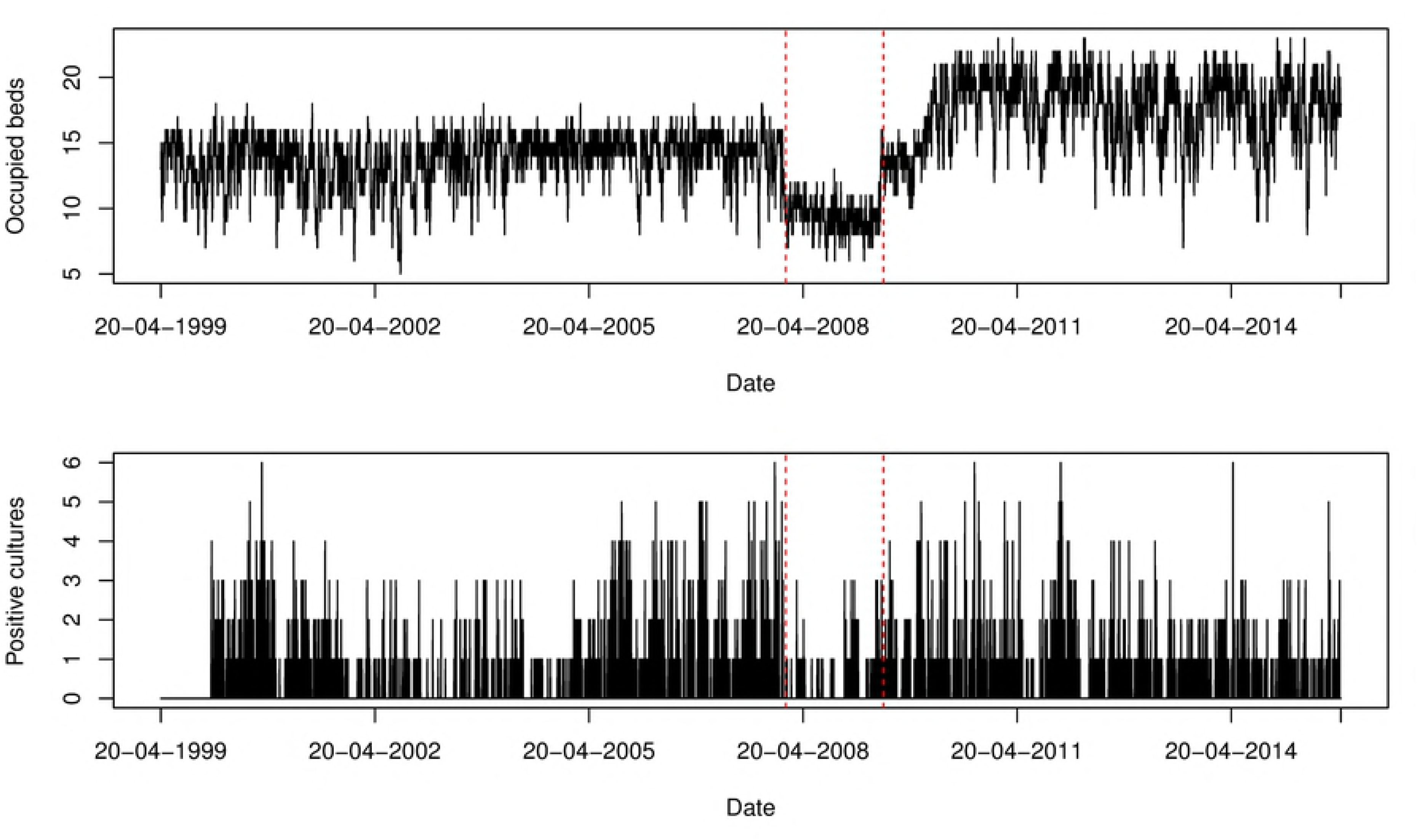
Number of occupied beds and positive isolates cultured from patients per swab day for ICU A. The red dotted lines indicate the time points that splits the study period into *before renovation* (20/04/1999 – 23/01/2008) and *after renovation* (06/11/2010 – 21/03/2017). Since the number of beds decreased during the renovation, the period is removed from the analysis.

**Fig 4.**
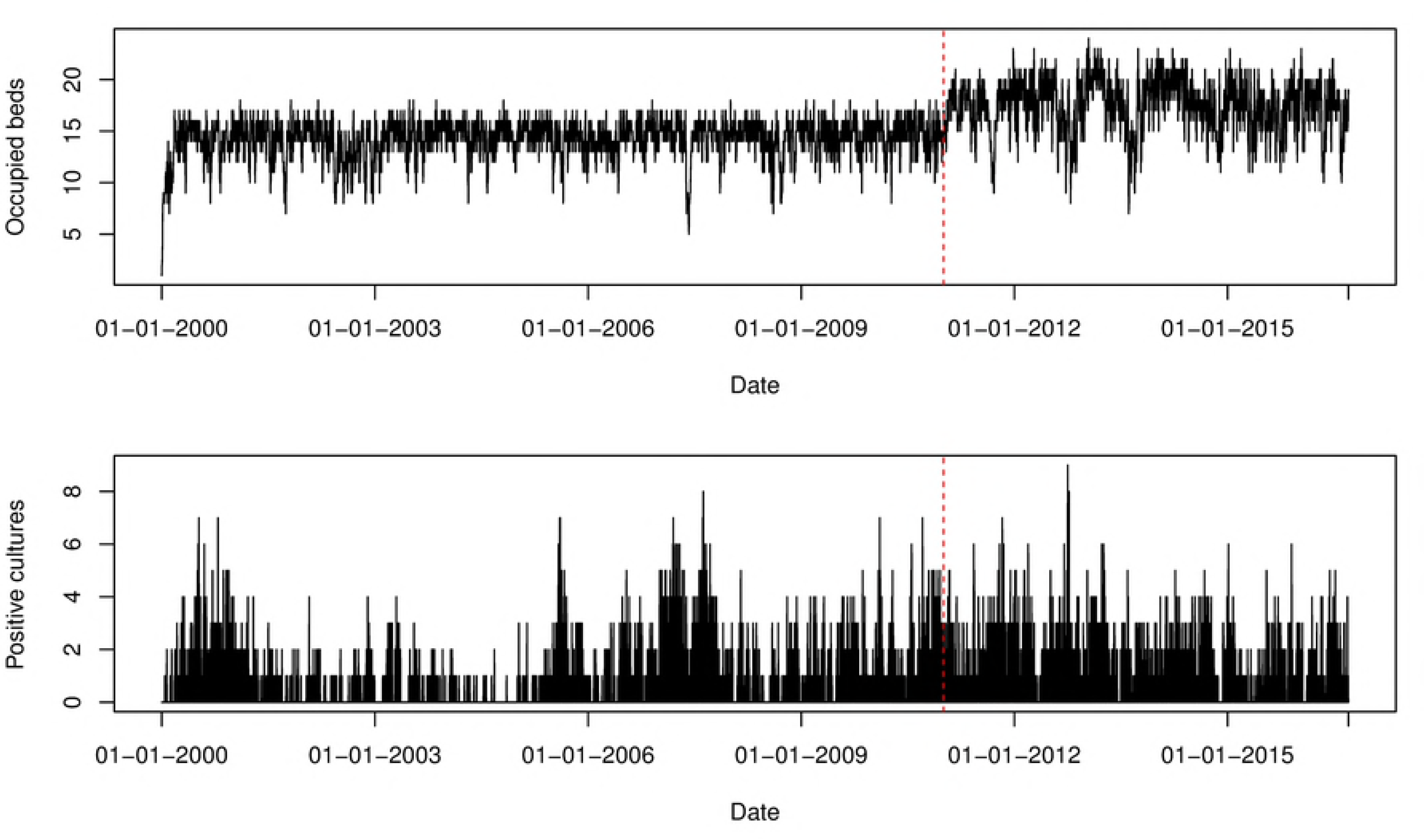
Number of occupied beds and positive isolates cultured from patients per swab day for ICU B. The red dotted line indicates the time point that splits the study period into *before renovation* (11/01/2000 – 12/01/2011) and *after renovation* (13/01/2011 – 13/09/2016).

### Estimated model parameters

Two model variants were fitted to the Besancon ICU data aiming to estimate the set of parameters *θ*_1_ = {*α*, *β*, *ϕ*, *f*} and *θ*_2_ = {*α*, *β*, *ε*, *μ*, *ϕ*, *f*} corresponding to the submodel with only two and the full model with all three transmission routes, respectively.

### Submodel: Two transmission routes

Posterior estimates of the model parameters for each ICU and each time period are reported in Table 2. Acceptance probabilities for proposed updates to the augmented data ranged from 3.2% (ICU B after renovation) to 11.1% (ICU A before renovation). Pairwise scatter plots indicated little correlation between parameter values, with the exception of a negative correlation between *α* and *β* (see S11). Histogram and trace plots of the posterior estimates are given in S12-S15 Figs and show that the MCMC chains rapidly mix and quickly converge to their stationary distribution. We found our estimates to be robust to the choice of priors for transmission parameters.

**Table 2.** Summary statistics of the marginal posterior distributions for model parameters.

| Parameter | Symbol | Median (95% credibility interval)* |  |  |  |
| --- | --- | --- | --- | --- | --- |
|  |  | ICU A |  | ICU B |  |
|  |  | BEFORE<sup>†</sup> | AFTER<sup>‡</sup> | BEFORE | AFTER |
| Endogenous coefficient | $\alpha$ | 0.011 (0.006, 0.016) | 0.013 (0.009, 0.016) | 0.007 (0.005, 0.01) | 0.014 (0.009, 0.018) |
| Cross-transmission coefficient | $\beta$ | 0.043 (0.021, 0.064) | 0.008 (0, 0.026) | 0.046 (0.032, 0.06) | 0.011 (0, 0.029) |
| Sensitivity | $\phi$ (%) | 50.9 (47.8, 54.2) | 50.2 (46.0, 54.4) | 61.8 (59.6, 64.0) | 58.6 (56.1, 61.1) |
| Importation probability | $f$ (%) | 4.5 (3.1, 6.1) | 6.2 (4.8, 7.8) | 6.0 (4.8, 7.1) | 9.9 (8.2, 11.6) |
| Fraction colonized | $p_{\text{col}}$ (%) | 24.4 (23.1, 25.8) | 19.9 (18.7, 21.2) | 22.3 (21.6, 23.0) | 24.4 (23.7, 25.1) |
| **Contributions** |  |  |  |  |  |
| Endogenous contribution | $R_{\text{end}}$ (%) | 53.6 (32.8, 75.9) | 89.3 (67.9, 100) | 43.4 (29.1, 58.7) | 84.5 (60.9, 100) |
| Cross-transmission contribution | $R_{\text{crossT}}$ (%) | 46.4 (24.1, 67.2) | 10.7 (0, 32.1) | 56.6 (41.3, 70.9) | 15.5 (0, 39.1) |

\*Highest posterior density interval
<sup>†</sup> 20/04/1999 – 23/01/2008 for ICU A and 11/01/2000 – 12/01/2011 ICU B.
<sup>‡</sup> 06/11/2010 – 21/03/2017 for ICU A and 13/01/2011 – 13/09/2016 for ICU B.

The probability of being colonized with *P. aeruginosa* on admission and the screening test sensitivity varied between the two ICUs and the time periods. For both ICUs, the median estimates of the importation probability *f* is higher in the data set after renovation than before, i.e. 4.5% and 6.2% for ICU A and 6.0% and 9.9% for ICU B. The difference between the time periods is only significant for ICU B. We estimated the median of the prevalence of *P.A*. to be 24.4% and 19.9% for ICU A and 22.3% and 24.4% for ICU B before and after renovation, respectively. Median estimates for the screening test sensitivity were 50.9% and 50.2% for ICU A and 61.8% and 58.6% for ICU B. Since the credibility intervals of the sensitivity estimates do not overlap with respect to the two ICUs, we can conclude that there is a 95% probability that the test sensitivity is higher in ICU B than in ICU A. Our possible explanation is based on the fact that the ICUs differ in their patient population. As a medical ward, ICU B contains patients with longer lengths of stay and more readmissions. Patients who are exposed to an ICU environment for a longer period of time may have a higher probability to get colonized at a detectable level. However, our explanation is only hypothetical and the true reason for the difference is not known.

The relative importance of the two considered transmission routes per ICU and time period is depicted in Fig 5 (a) and (b). For ICU A, the median relative contribution of the endogenous route is 53.6% (95% CrI: 32.8 – 75.9%) and 89.3% (95% CrI: 67.9 – 100%) leaving 46.4% (95% CrI: 24.1 – 67.2%) and 10.7 (95% CrI: 0 – 32.1%) of the acquisitions assigned to cross-transmission before and after renovation, respectively. For ICU B, 43.4% (95% CrI: 29.1 – 58.7%) and 84.5% (95% CrI: 60.9 – 100%) of the acquisitions were due to the endogenous route and cross-transmission accounted for 56.6% (95% CrI: 41.3 – 70.9%) and 15.5% (95% CrI: 0 – 39.1%) of the acquisitions before and after renovation, respectively.

**Fig 5.**
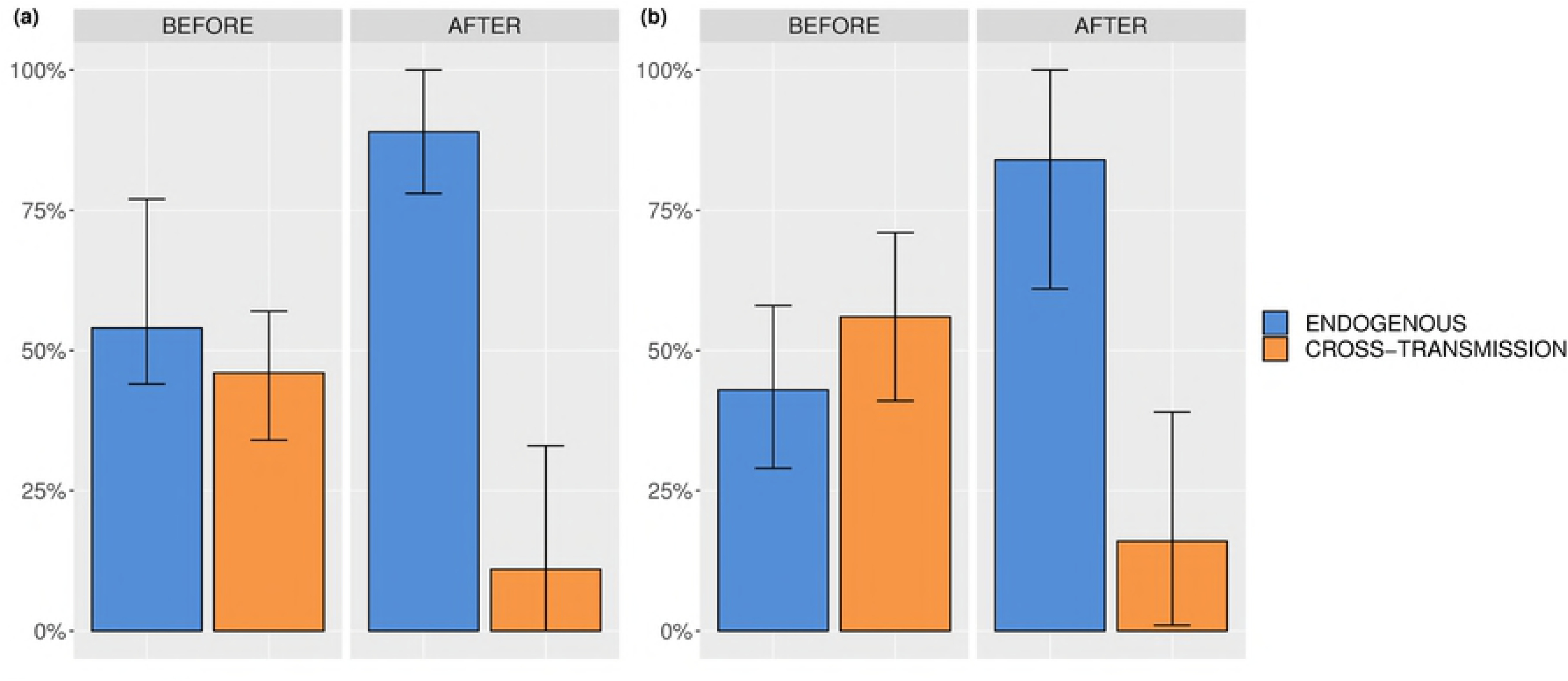
Relative contributions of endogenous route and cross-transmission. ICU A before and after renovation, (b) ICU B before and after renovation.

The results suggest that both routes have an important impact on the acquisitions in both ICUs. The median estimates of the relative contribution of the endogenous route are higher after than before renovation in both ICUs. Thus, there is a tendency for lower contribution of cross-transmission route after renovation in both ICUs. Possibly, hygiene was improved after renovating the ICUs. However, since the credibility interval for the endogenous route overlap before and after renovation, there is no evidence that the relative contributions differ between the time periods. Before renovation, the credibility intervals of the relative contributions for the endogenous route and cross-transmission overlap. Thus, we conclude that no route considerably predominates the transmissions before renovation. On the other hand, the respective credibility intervals do not overlap after renovation. Hence, the endogenous route predominates the transmissions after renovation. Comparing the results across ICUs, we can see that the credibility intervals of the relative contributions overlap leading to the conclusion that the two ICUs do not seem to be different regarding the relative importance of the transmission routes.

### Full model: Three transmission routes

Posterior estimates of the model parameters for each ICU are reported in Table 3. The estimates and interpretations for the importation rate *f*, the screening test sensitivity *ϕ* and the mean prevalence stay roughly the same when adding environmental contamination as an additional route. The same holds for the median relative contributions of the endogenous route and cross-transmission. The median relative contribution of environmental contamination after discharge is less than 1% ranging from 0.3% to 0.5% for both ICUs and both time periods. The relative importance of the three considered transmission routes per ICU and time period is depicted in Fig 6 (a) and (b). Acceptance probabilities for proposed updates to the augmented data ranged from 7.2% (ICU B after renovation) to 90% (ICU A before renovation). Pairwise scatter plots indicated strong correlations between *α* and *β*, *β* and *ε* and between *ε* and *μ* (see Fig 24). The correlation coefficient of the latter pair ranged from 0.531 to 0.561. Furthermore, it can be seen in Table 3 that the credibility intervals for the parameters *ε* and *μ* are large. Nevertheless, histogram and trace plots of the posterior estimates show that the MCMC chains rapidly mixed and quickly converged to their stationary distribution as can be seen in S16-S23 Figs. The rapid convergence could be achieved by tuning the parameters of the proposal distribution for *μ*. In contrast, a flat prior for the decay rate *μ* in combination with a small initial standard deviation for its proposal distribution resulted in large acceptance ratios close to 1. The MCMC chain mixed too slowly and therefore hindered the identifiability of the likelihood. This can be explained by the fact that our developed model is overparametrized when colonizations of patients are not or hardly influenced by environmental contamination. Small values of the transmission parameter *ε* as well as high values of the decay rate *μ* would reflect the aforementioned situation. As a result, the respective likelihood might not be or only weakly identifiable. Our sensitivity analyses and artificial data simulations demonstrated similar pairwise scatter plots and wide credibility intervals for the parameters *ε* and *μ* in case of a small contribution of environmental contamination to the transmissions (more details can be found in S7 Text). Hence, we can conclude that the role of environmental contamination after discharge within the transmission process of *P. aeruginosa* in the two ICUs A and B is small before as well as after renovation.

**Fig 6.**
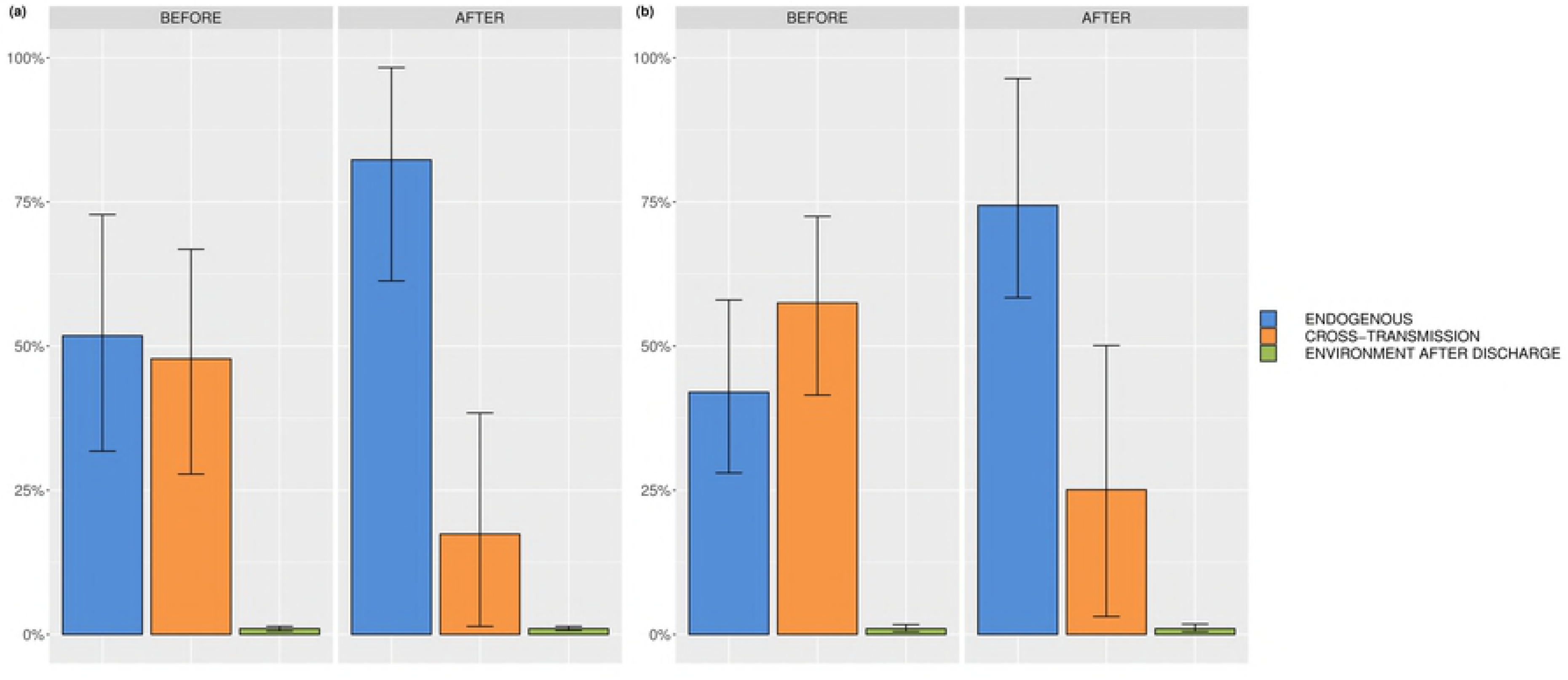
Relative contributions of endogenous route, cross-transmission and environmental contamination after discharge. (a) ICU A before and after renovation, (b) ICU B before and after renovation.

**Table 3.**
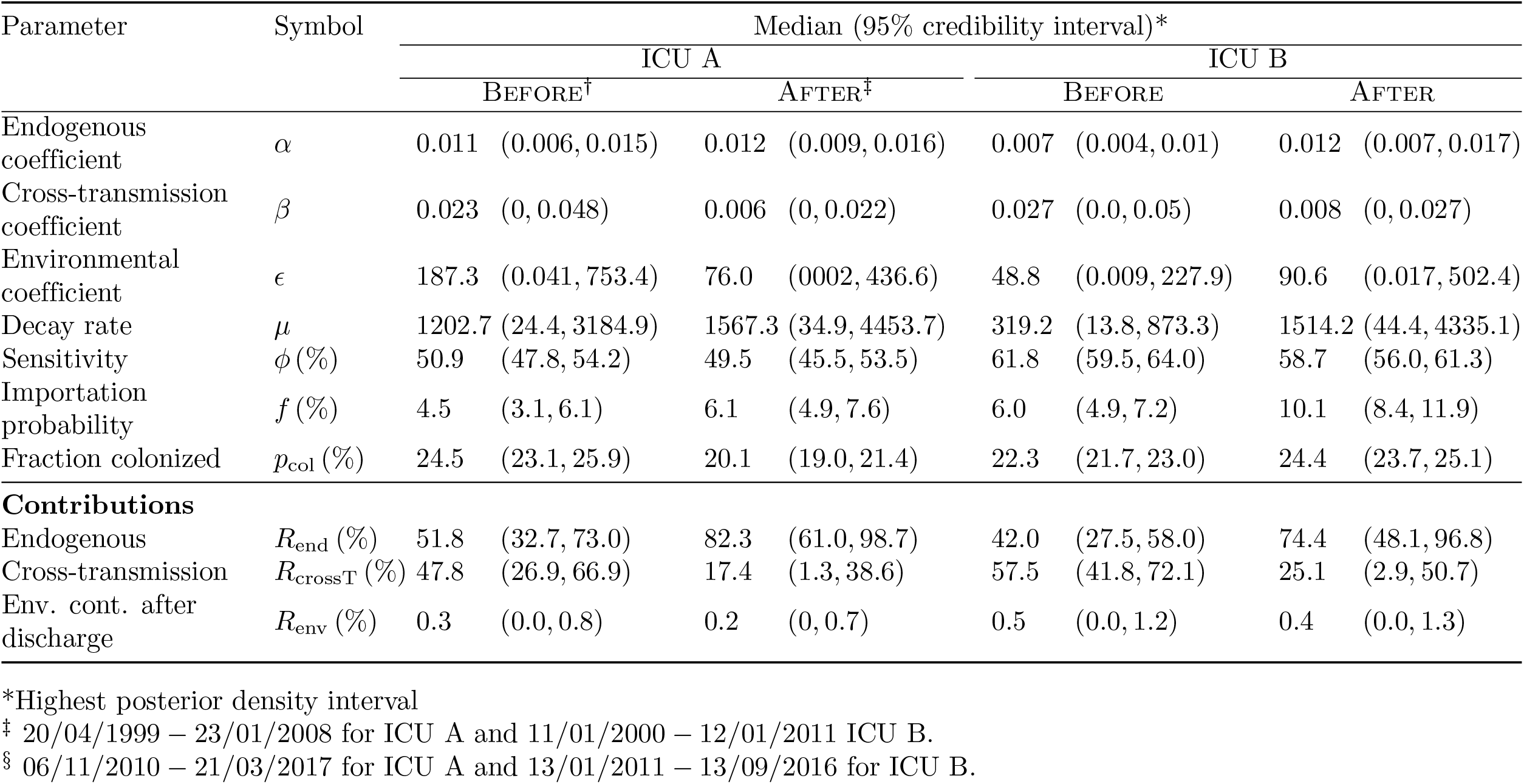
Summary statistics of the marginal posterior distributions for model parameters of the full model.

| Parameter | Symbol | Median (95% credibility interval)* |  |  |  |  |  |  |  |
| --- | --- | --- | --- | --- | --- | --- | --- | --- | --- |
|  |  | ICU A |  |  |  | ICU B |  |  |  |
|  |  | BEFORE<sup>†</sup> |  | AFTER<sup>‡</sup> |  | BEFORE |  | AFTER |  |
| Endogenous coefficient | $\alpha$ | 0.011 | (0.006, 0.015) | 0.012 | (0.009, 0.016) | 0.007 | (0.004, 0.01) | 0.012 | (0.007, 0.017) |
| Cross-transmission coefficient | $\beta$ | 0.023 | (0, 0.048) | 0.006 | (0, 0.022) | 0.027 | (0.0, 0.05) | 0.008 | (0, 0.027) |
| Environmental coefficient | $\epsilon$ | 187.3 | (0.041, 753.4) | 76.0 | (0002, 436.6) | 48.8 | (0.009, 227.9) | 90.6 | (0.017, 502.4) |
| Decay rate | $\mu$ | 1202.7 | (24.4, 3184.9) | 1567.3 | (34.9, 4453.7) | 319.2 | (13.8, 873.3) | 1514.2 | (44.4, 4335.1) |
| Sensitivity | $\phi$ (%) | 50.9 | (47.8, 54.2) | 49.5 | (45.5, 53.5) | 61.8 | (59.5, 64.0) | 58.7 | (56.0, 61.3) |
| Importation probability | $f$ (%) | 4.5 | (3.1, 6.1) | 6.1 | (4.9, 7.6) | 6.0 | (4.9, 7.2) | 10.1 | (8.4, 11.9) |
| Fraction colonized | $p_{\text{col}}$ (%) | 24.5 | (23.1, 25.9) | 20.1 | (19.0, 21.4) | 22.3 | (21.7, 23.0) | 24.4 | (23.7, 25.1) |
| **Contributions** |  |  |  |  |  |  |  |  |  |
| Endogenous | $R_{\text{end}}$ (%) | 51.8 | (32.7, 73.0) | 82.3 | (61.0, 98.7) | 42.0 | (27.5, 58.0) | 74.4 | (48.1, 96.8) |
| Cross-transmission | $R_{\text{crossT}}$ (%) | 47.8 | (26.9, 66.9) | 17.4 | (1.3, 38.6) | 57.5 | (41.8, 72.1) | 25.1 | (2.9, 50.7) |
| Env. cont. after discharge | $R_{\text{env}}$ (%) | 0.3 | (0.0, 0.8) | 0.2 | (0, 0.7) | 0.5 | (0.0, 1.2) | 0.4 | (0.0, 1.3) |

\*Highest posterior density interval
<sup>†</sup> 20/04/1999 – 23/01/2008 for ICU A and 11/01/2000 – 12/01/2011 ICU B.
<sup>‡</sup> 06/11/2010 – 21/03/2017 for ICU A and 13/01/2011 – 13/09/2016 for ICU B.

### Model selection

In total, 14 analyses were performed. For each ICU, three data sets were created - one for each time period and one combining the data sets before and after renovation. Additionally, the ICUs and time periods were combined in one data set. Each of the seven data sets were analyzed using the submodel and the full model. The DIC values for each model analysis can be found in Table 4. The analysis combining both ICUs and time periods shows smaller DIC values, i.e. 136507.8 and 130693, than the sum of the DICs for separate analyses (152428.8 and 150914.2) for both the submodel and full model, respectively. The full model results in a smaller DIC value for the analysis of the combined data set. Hence, based on the DIC, it would be sufficient to analyze the combined data set using the full model including endogenous route, cross-transmission and environmental contamination. Nevertheless, it can be seen in S8 Text that the posterior estimates of the different analyses are similar, especially for the relative contribution of environmental contamination after discharge.

**Table 4.**
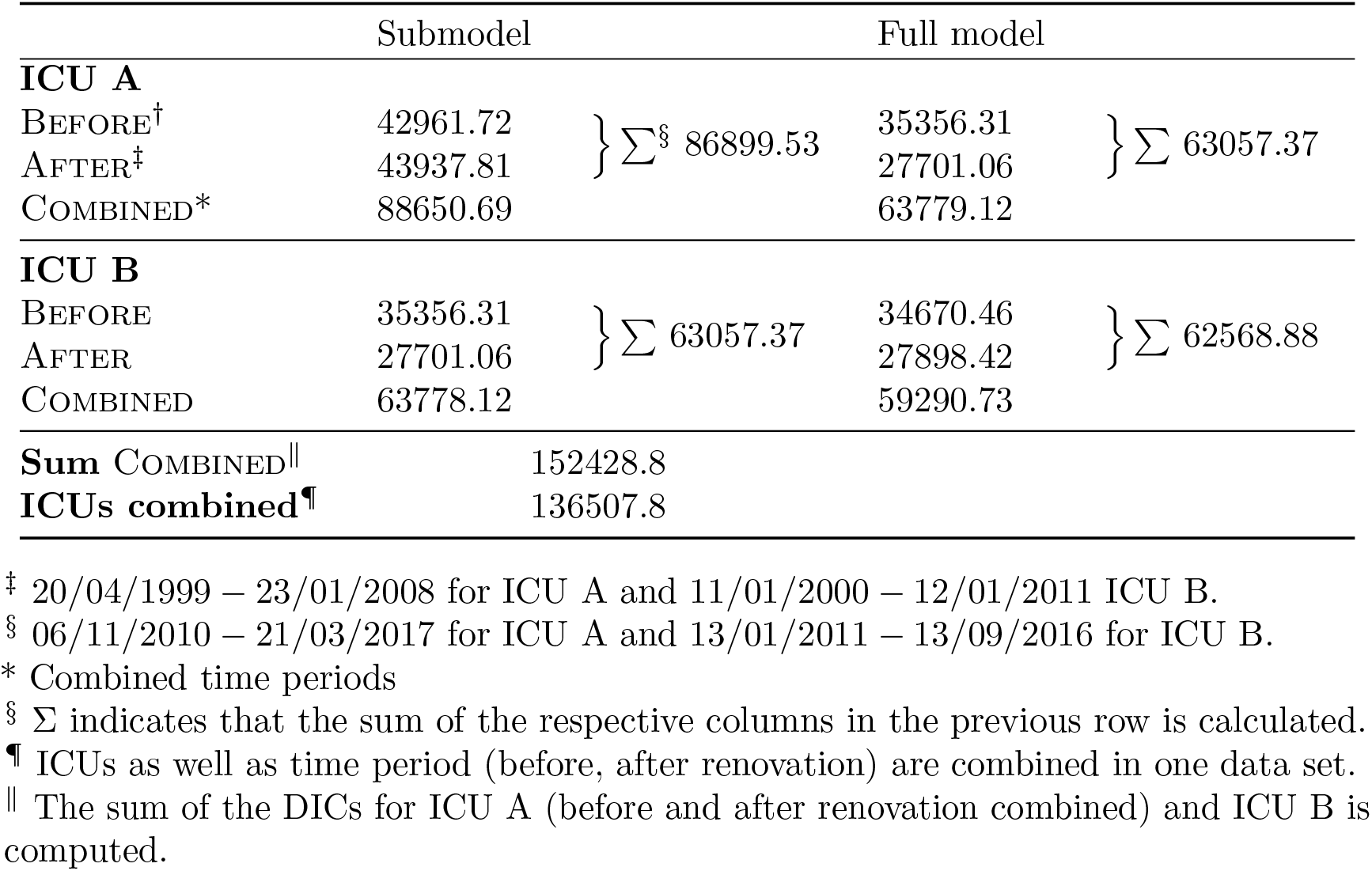
Deviance information criterion for the different models.

### Model assessment

For each bin of the force of infection the coverage probabilities are plotted and can be found in S25-S26 Figs. It can be seen that the coverage probabilities are approximately (sometimes higher, sometimes smaller) equal to the nominal confidence level of 0.95. Thus, both the full model as well as the submodel gave adequate fits to the four data sets. In Fig 7, the predicted fraction of acquisitions are plotted against the binned force of infection for one exemplary MCMC update. The red lines indicate the relationship between the probability of acquisition and force of infection assumed by our models. For this example, it is always contained by the confidence intervals (blue lines).

**Fig 7.**
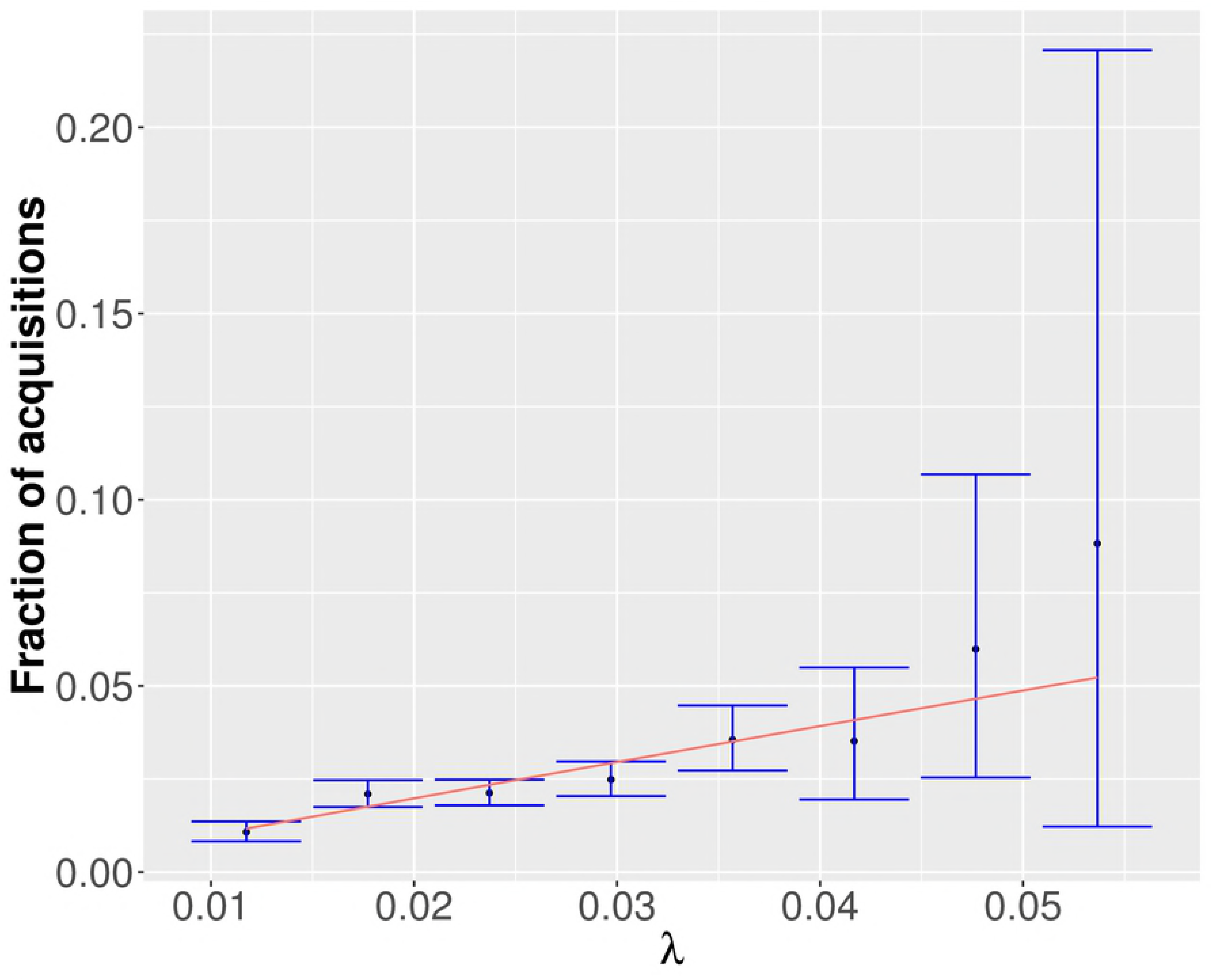
Exemplary model assessment plot for one MCMC update using the submodel applied to ICU A before renovation. The predicted fraction of acquisition is plotted against the theoretical force of infection. The red line indicates the theoretical relation between the force of infection and the probability of acquisition. The blue lines indicate 95% credibility intervals.

## Discussion

To our knowledge, our study is the first to use mathematical transmission models to estimate the relative contribution of environmental contamination after discharge for *P. aeruginosa* using only admission, discharge and screening data. The three different routes, endogenous route, cross-transmission and environmental contamination after discharge, are distinguished by the resulting patterns of the prevalence that they induce. We estimated that environmental contamination after discharge accounts for at most 1% of the total *P. aeruginosa* transmissions in the two ICUs of the University hospital in Besancon before and after renovation. In contrast, endogenous as well as cross-transmission are both essential for the transmission dynamics of *P. aeruginosa*. This suggests, that improvement of cleaning of the environment *after discharge* would have only a limited benefit regarding the prevention of *P. aeruginosa* colonization in the two considered ICUs of the University hospital in Besancon.

Previously, studies have been conducted to investigate the role of environmental contamination for colonizations of *P. aeruginosa*,. For instance, Panagea et al. performed environmental studies to determine the extent of environmental contamination with an epidemic strain of *P. aeruginosa* [25]. They concluded that the transmissibility of the epidemic strain cannot be explained solely on the basis of improved environmental survival. Our results likewise demonstrate that the decay of *P. aeruginosa* is already rapid enough to limit its survival in the environment.

While our approach is efficient in determining the relative contribution of environmental contamination after discharge requiring merely longitudinal surveillance data, it has several limitations that may restrict its practical applicability.

Our conclusions on the impact of cleaning only applies to the environment after the discharge of patients. Permanently contaminated reservoirs in ICUs, such as sinks, may still serve as sources for colonization. In our model they are assigned to the endogenous route. Thus, while the effect of cleaning improvement after discharge might be limited for the two considered ICUs, general cleaning improvement of the environment might be important to reduce permanent reservoirs for environmental contamination.

The results of our analysis build on a data-augmented MCMC algorithm [19,26]. Markov chain Monte Carlo sampling is a powerful tool to estimate posterior parameter distribution whenever the likelihood is analytically intractable. And yet, the inherent disadvantage of this sampling scheme is that it may take prohibitively many iterations to converge to the posterior distribution. The convergence properties of MCMC sampling in high-dimensional posterior distributions can be particularly problematic and sensitive to the choice of prior and proposal distributions. Thus, tuning of the MCMC parameters becomes crucial for its application. Our developed full model is overparametrized when colonizations of patients are not or hardly influenced by environmental contamination. As a result, the respective likelihood might not be identifiable or only weakly identifiable. Here, a flat prior for the decay rate *μ* in combination with a small initial standard deviation for its proposal distribution resulted in large acceptance ratios close to 1. The MCMC chain mixed too slowly and therefore hindered the identifiability of the likelihood. We were able to tune the parameters of the proposal distribution for *μ* such that rapid convergence to the posterior distribution could be assessed using visual inspection of histograms and trace plots. However, as presented in the *Results* section, pairwise scatter plots showed strong correlations in particular between *ε* and *μ*. Simulation studies confirmed that this can be explained by an absence of environmental contamination in the investigated data sets. This supports our finding that an impact of environmental contamination after discharge on the transmission of *P. aeruginosa* may be neglected.

Moreover, colonization is assumed to remain until discharge. While this assumption is true for *P. aeurginosa* it does not hold true for all antibiotic-resistant nosocomial pathogens. However, intermittent carriage may be readily included allowing the method to be generalized to other pathogens.

We assumed no difference in transmissibility between different strains of *P. aeruginosa* and that all colonized patients are equally likely to transmit the pathogen. While information on antibiotic resistance or microbial genotyping in combination with epidemiological data may aid in distinguishing different strains and identifying specific transmission events, only the uncertainty of the estimates would be affected. In particular, the widths of the credibility intervals are likely to be reduced, but we do not expect a large effect on the parameter estimates.

Assessing the fit of the model to the data is crucial to model building. The true relative importance of the different routes of colonization in ICUs is generally unknown. Genotyping data that might be used to demonstrate the source of the acquired colonization is generally scarce and was not available for the data used in our analysis. While the posterior predictive *p*-value is a popular method for assessing model fit, it has been increasingly criticized for its self-fulfilling nature [27]. Furthermore, the choice of the test statistic is crucial in order to adequately summarize discrepancies between datasets. Rather than relying on a suitable summary statistic, we presented a model assessment method that evaluates whether the estimated force of infection adequately represents the transmission dynamics in the ward. However, while the corresponding coverage probabilities may depict discrepancies per bin of the force of infection, the sample size is not controlled by choosing the number MCMC updates. It might well occur that specific patients (and their acquisition events) appear in more than one MCMC update simultaneously. Thus, the true sample size is estimated to be smaller. Further improvement of the method presented here or development of other methods would be a vital topic for assessing epidemic models.

Model selection was performed using the DIC which is known to display poor performance (i.e. identifying the correct model) for complex likelihood functions such as those corresponding to epidemic models. Comparing the plausibility of different models is crucial for selecting the model that describes the dynamics of the observed system best. Nevertheless, model choice for stochastic epidemic models is far from trivial. All known approaches for model selection exhibit advantages as well as disadvantages [27] which makes selecting the most suitable model comparison technique not straightforward. We selected the well-known DIC-method that was easy to use and implement. Our main results regarding environmental contamination after discharge do not depend on the model choice. And yet, the development of a suitable and robust model selection procedure in a data-augmented Bayesian framework would be an interesting and important topic for future research.

Finally, like all models, ours is a simplification of the truth as it is unlikely that all relevant variables are already included. Adding covariates such as antibiotic use, sex or age may improve the model fit.

Our work may be used or further extended for assessing the relative importance of different transmission routes within intensive-care units not only for *P. aeruginosa* but for hospital pathogens in general. Based on these results, consequential decisions for tailored interventions or policies may be deduced, aiding in improving infection prevention and control and therefore reducing morbidity, mortality and related costs in hospitals.

## Acknowledgments

The research leading to these results was conducted as part of the COMBACTE-MAGNET (Combatting Bacterial Resistance in Europe - Molecules Against Gram-Negative Infections) consortium. For further information please refer to www.COMBACTE.com.

## Supporting information

### S1 Text. Environmental contamination

The full model includes environmental contamination on a ward-level. The bacterial load at any given time *t* is based on the differential equation

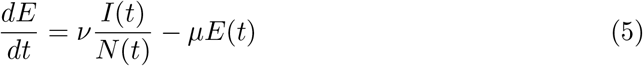

Solving the above differential equation requires discretizing over *t*, resulting in a finite number of time steps *t*_0_, *t*_1_, …, *t*_*N*_. We then assume *I*(*t*) = *I_t_* and *N*(*t*) = *N_t_* to be constant within a time step and use it as initial conditions. Separating variables leads to

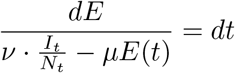

and thus

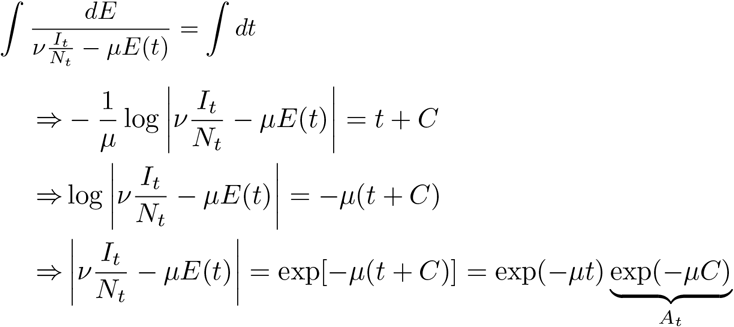

Now, two cases have to be distinguished.

1. Case: 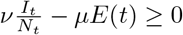

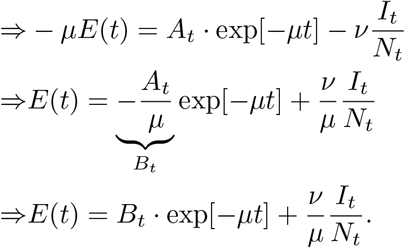
2. Case: 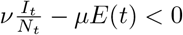

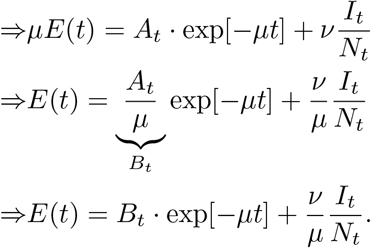

Determine 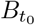 for initial condition 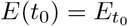:

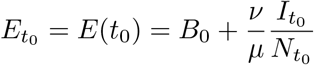

and therefore

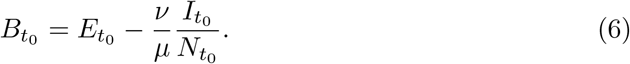

For *t*_0_ ≤ *t* ≤ *t*_1_ the environmental load can be then computed by

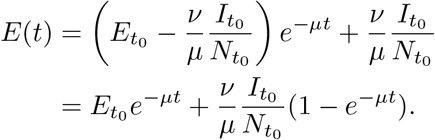

For *t*_0_ ≤ *t*_*i*_ ≤ *t_N_* it holds

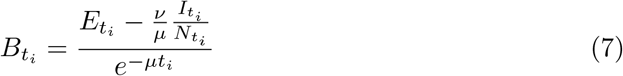

and therefore, it holds for ⎿*t*⏌:= max{*t*_0_ ≤ *x* ≤ *t_N_* | *x* ≤ *t*} and *t* ∈ ℝ \ {*t*_0_, *t*_1_,…, *t_N_*}

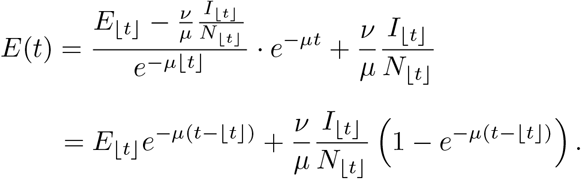

and

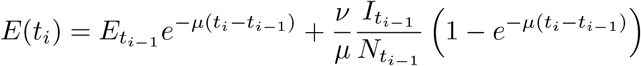

for 0 ≤ *i* ≤ *N*.

### S2 Text. Discrete-time transmission model

For the discrete-time transmission model, we assume that the number of colonized patients *I*(*t*), the total number of patients *N*(*t*) and the bacterial load *E*(*t*) is constant during the day. It is assumed that admission and screening occur at 12:00 pm on each day *T* determining *I*_*T*_ and *N*_*T*_. Given all the information (at 12:00 pm), the environmental contamination on day *T* is determined. The force of infection on day *T* is then given by

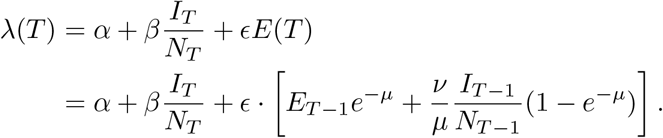

### S3 Text. Relative contribution

The computations in section *Relative contributions of transmission routes* were developed for continuous-time models. In our discrete-time model, we assume that events such as, admission, colonization and discharge of patients and screening occur on a daily basis. However, we do assume that the level of environmental contamination changes continuously. Computing the relative contributions of the different transmission routes becomes more laborious in this scenario. Let 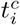 be the acquisition time of patient *i* ∈ {1,…, *n*}. The contribution of a route *j* is the ratio of the probability that the acquisition was due to route *j* and the total probability of acquisition:

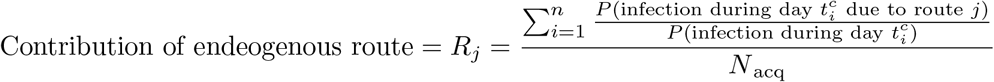

where *N*_acq_ is the total number of occured colonizations and *R*_*j*_ with *j* ∈ {end, crossT, env} indicate the endogenous, cross-transmission or environmental route, respectively. The route-specific probabilities can be determined by

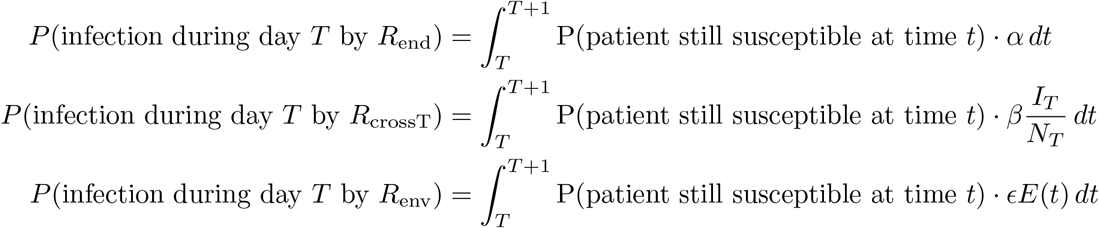

where environmental contamination during a patient’s stay is assigned to the environmental route. For the alternative assignment (i.e. only considering the bacterial load remaining after discharge as environmental contamination), the formulas change according to Eq. (3). The results are dependent on 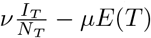 and are given by

1. Case: 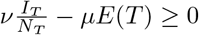

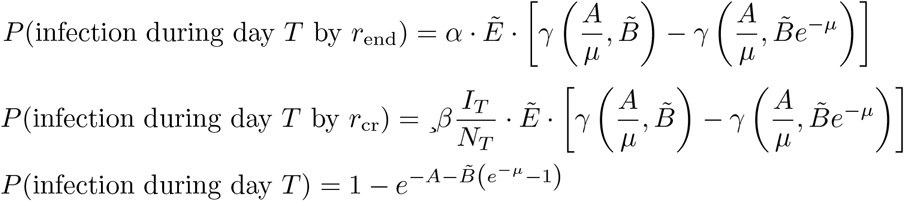
2. Case: 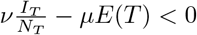

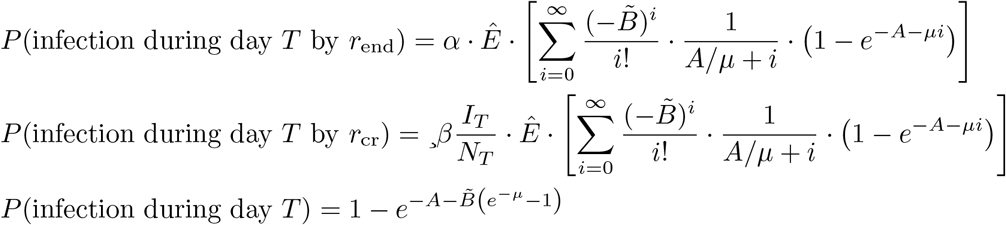

and

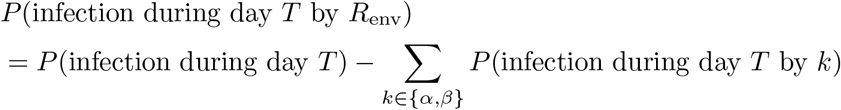

where

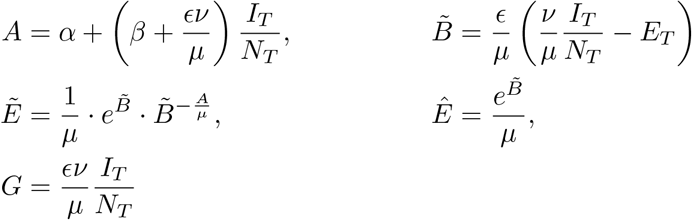

and *γ*(·,·) is the lower incomplete gamma function. Note that the derivations are omitted here but can be requested from the first author.

### S4 Text. Approximation of relative contribution in discrete-time

Large values of the force of infection *λ*(*t*) are very unlikely. Under the assumption of small *λ*(*t*), the following simplifications and approximations can be made:

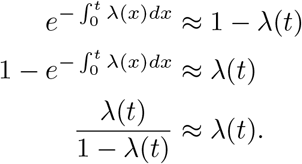

Therefore, the force of infection itself may be a good approximation of the probability of infection and the probability of acquiring colonization due to route *j* may be approximated by the respective sub-term of the force of infection assigned to route *j*:

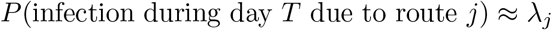

with *j* ∈ {end, crossT, env}. As an approximation of the relative contribution we compute the ratio of the transmission rate and the force of infection for each acquired colonization:

- Contribution of endogenous route = 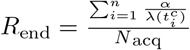
- Contribution of cross-transmission = 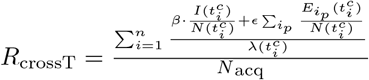
- Contribution of environmental contamination = 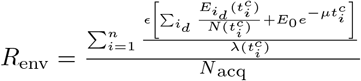

where 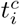 is the day of colonization of patient *i* ∈ {1,…, *n*} and *N*_acq_ the total number of occured colonizations. Furthermore, *i*_*p*_ indicates a colonized patient that is present at time 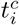 and *i*_*d*_ a colonized patient that has been colonized prior to 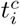 but was already discharged. It holds *R*_end_ + *R*_crossT_ + *R*_env_ = 1.

### S5 Text. Adapted data-augmented MCMC algorithm

We model the transmission process using a two-state Markov model, where each patient can be either *susceptible* (P.A. negative) or *colonized* (P.A. positive). A patient i is admitted to the ICU on day 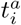 and discharged on day 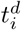. The probability that a patient is admitted already colonized is described by parameter *f*. The rate at which a susceptible patient transitions to being colonized is described in section *Materials and methods*. The colonization state of an individual patient is determined from screening information. We suppose that for each patient *i* a set of screening results 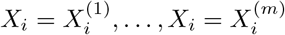, taken on days 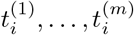 is available. The set of all screening results is denoted by *X* = {*X*_1_,…,*X*_*N*_} where *N* is the total number of patients. Since screening tests are typically intermittent and imperfect, we define the test sensitivity *ϕ* (i.e. probability that a colonized patient has a positive result). We assume that the specificity (i.e. probability that an uncolonized patient has a negative result) is 100%.

We implemented an adapted version of the data-augmented MCMC algorithm to analyze the data. The transmission and importation model, as well as the data-augmentation method is closely based on the approach of [19, 26] but adapted for the transmission routes presented in this paper. The algorithm was implemented in C++ and the analysis of the output was performed in `R` (Version 3.5.1) [28]. The aim of our analysis was to estimate the set of parameters *θ* = {*α*, *β*, *ε*, *μ*, *f*, *ϕ*}. The prior distribution were chosen as follows:

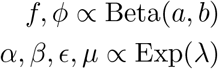

where Exp(*λ*) represents the exponential distribution with rate *λ*, and Beta(*a*, *b*) the beta distribution with shape parameters *a* and *b*. Having fixed *a* = *b* =1 and *λ* = 0.001, we use uninformative priors in our analysis.

The data-augmentation procedure accounts for unobserved colonization times by augmenting the parameter space with *A* = {*t^c^*, *s^a^*}, a set comprising of the unobserved colonization times *t*^*c*^ and admission states *s*_*a*_ of all *n* patients. An admission state of a patient is 1 if the patient is colonized upon admission and 0 otherwise. If the patient *j* becomes colonized during his/her stay, the colonization time may take an integer value between the time of admission 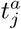 and time of discharge 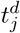 (inclusive). If a patient does not acquire colonization, the respective value 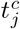 takes a dummy value of –1. The augmented posterior density relation can be determined using Bayes’ Theorem:

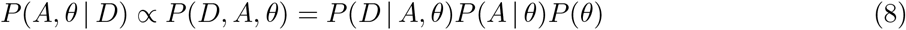

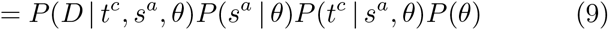

where *P*(*D* | *A*, *θ*) is the likelihood of the observed data *D*, *P*(*A* | *θ*) is the likelihood of the augmented data and *P*(*θ*) is the joint prior distribution of the parameter set *θ*. All terms in (8) can be explicitly calculated. It holds

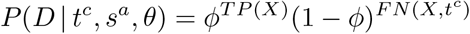

where *TP*(*X*) and *FN*(*X*, *A*) are the total number of true positive and false negative swab results, given the colonization times *t*^*c*^, respectively. It represents the imperfect observation of the transmission dynamics. Assuming that lost colonization can be excluded, we consider any negative result after the time of colonization as a false negative. Since false positive results are impossible, the *TP*(*X*) is not dependent on the augmented data and can be determined directly from the observed data. The probability of the set of importations, given the importation probability *f* is given by

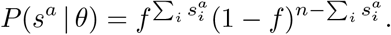

The transmission model itself is reflected in the probability of the colonization times given the admission states and the parameters

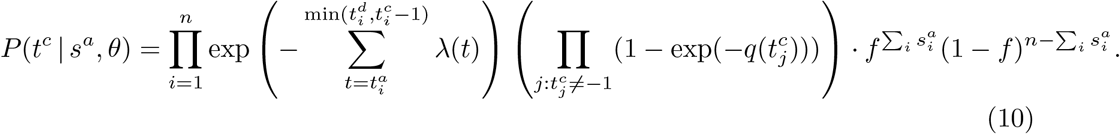

To update the importation rate *f* and the sensitivity *ϕ*, we use Gibbs sampling as we can sample directly from the full conditional distributions. The transmission parameters *α*, *β*, *ε*, *μ* are updated using an adapted version of the Metropolis-Hastings algorithm. Regular MCMC methods based on the Metropolis-Hastings algorithm tend to be very slow in high dimensions as a result of slow mixing and therefore inefficient convergence towards the target distribution. In high-dimensional spaces the volume outside is much larger than the volume of our target distribution. Thus, traditional MCMC methods such as the Metropolis-Hastings algorithm, spend considerable amount of time of traversing space away from the mode of the target distribution. Our adapted MCMC algorithm aims in exploring the target distribution more efficiently.

The Metropolis-Hastings algorithm generates a Markov chain *θ*^(1)^,…, *θ*^(*Ν*)^ which converges to a target distribution *π*(·) if *N* is large enough. In each update of the Markov chain, a candidate point, *θ*^*^ is sampled from a proposal density *q*(*θ*^*^ | *θ*^(*i*)^), which gives the probability density of proposing *θ*^*^, given the current, *i*^th^ value. With a certain probability or so-called acceptance ratio

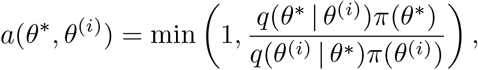

the proposed value is accepted.

The Metropolis algorithm is a special case of the Metropolis-Hastings algorithm where the proposal function is symmetrical. Since a symmetrical proposal distribution simplifies the calculation of the acceptance ratio to *a*(*θ*^*^, *θ*^(*i*)^) = min (1, *π*(*θ*^*^)/*π*(*θ*^(*i*)^)), it is often used for updating parameters. The proposal function has a great influence on the speed of convergence and hence efficiency of the algorithm. We suggest a proposal distribution that speeds up the convergence towards the target distribution while limiting the additional computational effort. The idea behind our method is as follows: For each estimated parameter set *θ* there is a corresponding force of infection *λ*(*t*) for each time *t*. It can be assumed that the mean force of infection 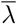 is approximately constant over the number of iterations. The rationale behind it is that there is *true* mean force of infection that should be approximated by the MCMC algorithm. Proposing new parameter candidates depending on the mean force of infection reduces the volume that has to be traversed in order to converge to the target distribution. The resulting proposal density is not symmetric anymore and thus the procedure requires an adjustment of the acceptance ratio. The adapted Metropolis-Hastings algorithm to update the transmission parameters runs as follows:

<u>Two transmission routes</u>

1. Set initial values *θ*^(0)^ = (*α*^(0)^, *β*^(0)^), and the number of iterations *N*.
2. Sample new parameter values *α*^*^,*β*^*^ as follows:

a. Propose candidate *α*^*^ by sampling from 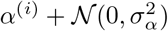
b. Propose candidate *β*^*^ assuming 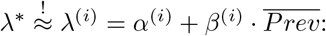: Sample *β*^*^ from 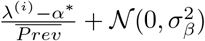, i.e. 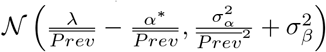
c. With probability

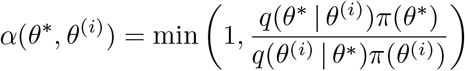

where 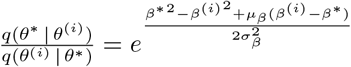, accept the proposed value and set *θ*^(*i*+1)^ = *θ*^*^, else set *θ*^(*i*+1)^ = *θ*^(*i*)^.
3. If *i* < *N*, then go to step 2.

<u>Three transmission routes</u>

1. Set initial values *θ*^(0)^ = (*α*^(0)^, *β*^(0)^, *ε*^(0)^, *μ*^(0)^), and the number of iterations *N*.
2. Sample new parameters *θ*^*^ = (*α*^*^, *β*^*^, *ε*^*^, *μ*^*^) from a proposal density *q*(*θ*^*^ | *θ*^(*i*)^) as follows:

a. Propose candidate *α*^*^ by sampling from 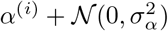
b. Propose candidate *β*^*^ by sampling from 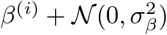
c. Propose candidate *μ*_1_ by sampling from 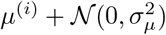
d. Update 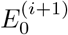 to 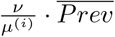
e. Propose candidate *ε*^*^ assuming 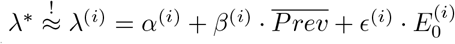: Sample *ε*^*^ from 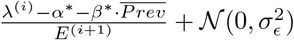, i.e. 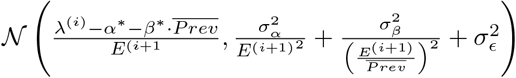.
f. With probability

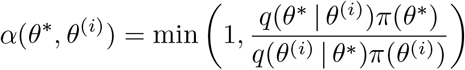

where 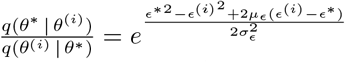, accept the proposed value and set *θ*^(*i*+1)^ = *θ*^*^, else set *θ*^(*i*+1)^ = *θ*^(*i*)^.
3. If *i* < *N*, then go to step 2.

### S6 Text. Model selection

We would like to assess whether we can concatenate the Besancon data e.g. before and after the renovation of the ICUs in one large data set to increase the power of our method. The idea is to compare the DICs for two different scenarios:

- Consider only one model including one parameter set *θ* = {*α*, *β*, *f*, *ϕ*} where *α* is the endogenous, *β* the cross-transmission parameter, *f* the importation rate and *ϕ* the test sensitvity. The analysis is then performed on all the data *X* of the two ICUs and the two time periods (before and after renovation). The DIC is then given by

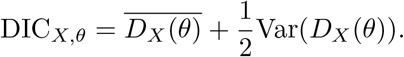
- Consider a model including a parameter set consisting of separate parameters for each time period:

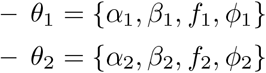 The parameter set of the model is then:

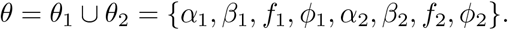 The parameters in *θ*_1_ are updated for the data set before renovation whereas the parameters in *θ*_2_ are updated for the data set after renovation. The deviance for this model is determined by

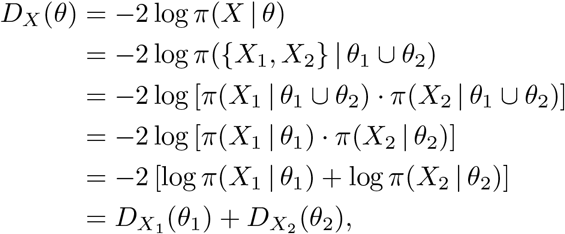

where *X*_1_ is the data set for the time period before and *X*_2_ for the time period after renovation. Thus, the DIC for the model including separate parameters for each time period can be calculated as

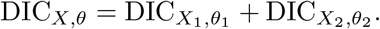

### S7 Text. Simulation studies

Several data sets were simulated to test the algorithm’s ability to identify the correct relative contribution. We fixed values for *α*, *β* and *ν* and varied values for *μ*. The parameter *ε* is set to 0.6 · *μ*. Three main scenarios regarding the duration of persistence of bacterial load in the environment are analyzed:

1. Long: *μ* ∈ {1/7,1/14}
2. Medium: *μ* = 1/3
3. Short: *μ* = 1

For each of the three scenarios the importation rate is varied within {0.01, 0.05, 0.1}. We observed that for long and short durations of bacterial persistence, chain convergence cannot be attained in a reasonable amount of time using uninformative exponential priors. A long duration of bacterial persistence leads to a model that cannot be distinguished from one with a higher endogenous contribution and small contribution of environmental contamination. On the other hand, a short duration of bacterial persistence leads to difficulties in distinguishing the resulting model from one with a higher contribution of cross-transmission and smaller contribution of environmental contamination. However, informative priors on *μ* (e.g. uniform priors *U*(0, 2)) led to more rapid convergence of the MCMC chain.

For a medium length of bacterial persistence, the model is able to estimate the simulated parameter values i.e. the true parameter values and respective relative contributions lie in the 95% credibility intervals, given the mean prevalence was large enough (> 15%). We have performed further simulation studies where the relative contribution of environmental contamination after discharge varied. In particular, when environmental contamination after discharge was not present in the simulated data, the results resembled our analysis of the Besancon data shown in the *Results* section. Further information on our simulation studies may be requested from the first author.

### S8 Text. Secondary analyses

In addition to the analyses presented in *Results*, six further analyses were performed. For each ICU, the time periods before and after renovation were combined. Finally, all available data was concatenated into one big data set and analyzed at once. The results of these analyses using the submodel as well as the full model are presented in S9 - S10 Tables. The posterior estimates of the model parameters and the corresponding relative contributions are similar to the ones presented in the *Results* section.

**S9 Table.**
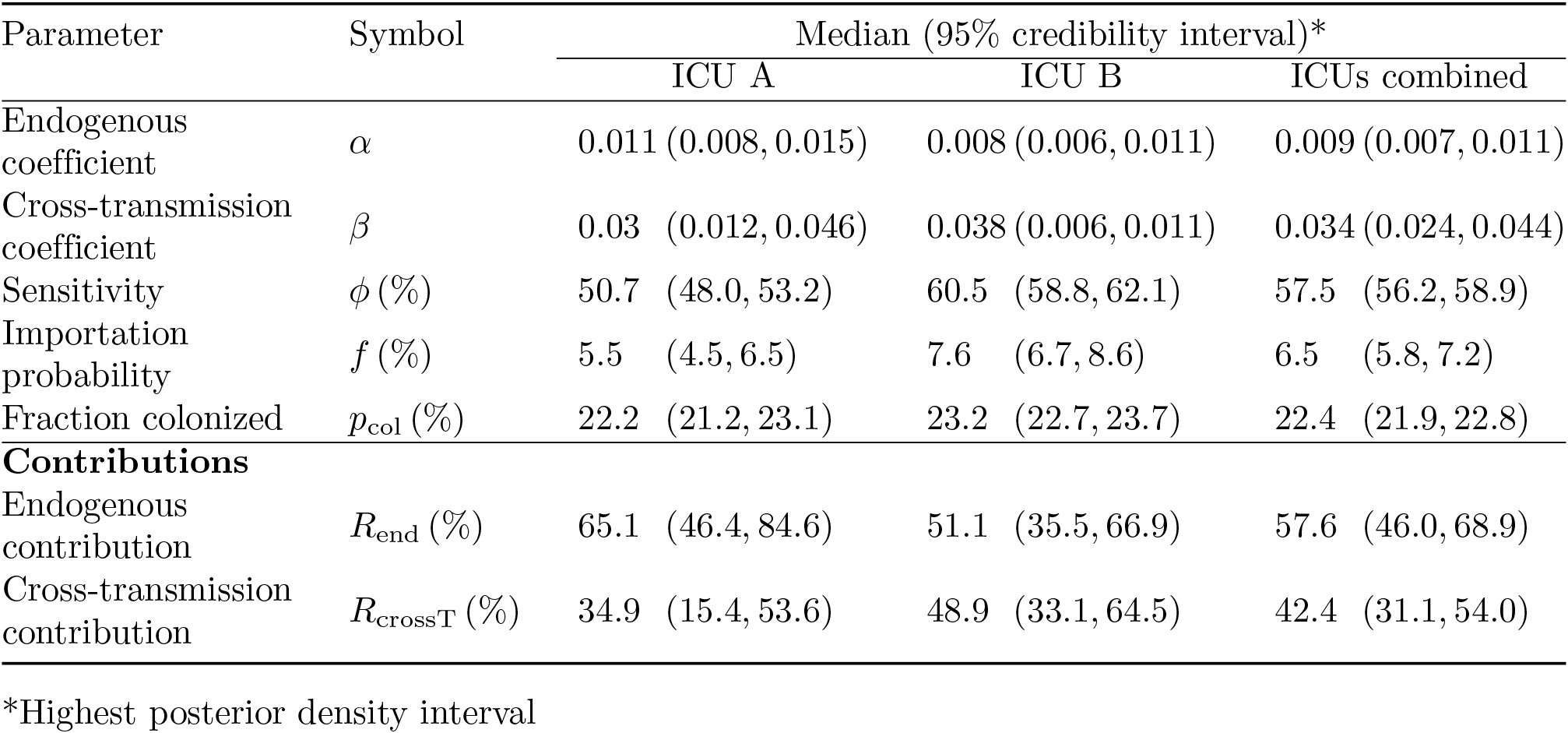
Summary statistics of the marginal posterior distributions for model parameters of the submodel.

**S10 Table.** Summary Statistics of the marginal posterior distributions for model parameters of the full model.

| Parameter | Symbol | Median (95% credibility interval)* |  |  |  |  |  |
| --- | --- | --- | --- | --- | --- | --- | --- |
|  |  | ICU A |  | ICU B |  | ICUs combined |  |
| Endogenous coefficient | $\alpha$ | 0.011 | (0.008, 0.014) | 0.008 | (0.006, 0.011) | 0.009 | (0.007, 0.011) |
| Cross-transmission coefficient | $\beta$ | 0.013 | (0, 0.033) | 0.022 | (0, 0.011) | 0.018 | (0, 0.034) |
| Environmental coefficient | $\epsilon$ | 201.1 | (0.018, 832.829) | 176.7 | (0.009, 811.6) | 209.0 | (0.02, 784.5) |
| Decay rate | $\mu$ | 1415.8 | (65.6, 4273.8) | 1396.9 | (26.4, 3992.2) | 1419.2 | (43.7, 4524.0) |
| Sensitivity | $\phi$ (%) | 50.6 | (48.0, 53.1) | 60.5 | (58.9, 62.1) | 57.6 | (56.2, 58.9) |
| Importation probability | $f$ (%) | 5.5 | (4.5, 6.6) | 7.5 | (6.6, 8.4) | 6.4 | (5.7, 7.2) |
| Fraction colonized | $p_{\text{col}}$ (%) | 22.2 | (21.3, 23.1) | 23.2 | (22.7, 23.6) | 22.4 | (21.9, 22.8) |
| **Contributions** |  |  |  |  |  |  |  |
| Endogenous | $R_{\text{end}}$ (%) | 62.2 | (44.8, 79.7) | 50.4 | (35.6, 65.2) | 57.6 | (45.5, 67.9) |
| Cross-transmission | $R_{\text{crossT}}$ (%) | 37.2 | (20.4, 54.9) | 49.0 | (34.0, 63.1) | 42.7 | (31.9, 54.0) |
| Env. cont. after discharge | $R_{\text{env}}$ (%) | 0.006 | (0, 0.013) | 0.005 | (0, 0.013) | 0.006 | (0, 0.012) |

\*Highest posterior density interval

**S11 Fig. Pairwise plots of samples from the posterior distribution for the transmission parameters of the submodel.** The plots were generated from the data of ICU A before renovation using the submodel with endogenous and exogenous transmission.

S12 Fig. Histograms for ICU A before renovation using the submodel with endogenous route and cross-transmission.

S13 Fig. Traceplots for ICU A before renovation using the submodel with endogenous route and cross-transmission.

S14 Fig. Histograms for ICU A after renovation using the submodel with endogenous route and cross-transmission.

S15 Fig. Traceplots for ICU A after renovation using the submodel with endogenous route and cross-transmission.

**S16 Fig. Histograms for ICU A before renovation using the full model with endogenous route, cross-transmission and environmental contamination.** The results are displayed for transmission parameters *α*, *β*, *ε* and *μ*.

**S17 Fig. Histograms for ICU A before renovation using the full model with endogenous route, cross-transmission and environmental contamination.** The results are displayed for importation probability *f*, sensitivity parameter *ϕ*, log-likelihood and relative contributions *R*_*i*_ where *i* ∈ {endogenous, cross-transmission, environment}.

**S18 Fig. Traceplots for ICU A before renovation using the full model with endogenous route, cross-transmission and environmental contamination.** The results are displayed for transmission parameters *α*, *β*, *ε* and *μ*.

**S19 Fig. Traceplots for ICU A before renovation using the full model with endogenous route, cross-transmission and environmental contamination.** The results are displayed for importation probability *f*, sensitivity parameter *ϕ*, log-likelihood and relative contributions *R*_*i*_ where *i* ∈ {endogenous, cross-transmission, environment}.

**S20 Fig. Histograms for ICU A after renovation using the full model with endogenous route, cross-transmission and environmental contamination.** The results are displayed for transmission parameters *α*, *β*, *ε* and *μ*.

**S21 Fig. Histograms for ICU A after renovation using the full model with endogenous route, cross-transmission and environmental contamination.** The results are displayed for importation probability *f*, sensitivity parameter *ϕ*, log-likelihood and relative contributions *R*_*i*_ where *i* ∈ {endogenous, cross-transmission, environment}.

**S22 Fig. Traceplots for ICU A after renovation using the full model with endogenous route, cross-transmission and environmental contamination.** The results are displayed for transmission parameters *α*, *β*, *ε* and *μ*.

**S23 Fig. Traceplots for ICU A after renovation using the full model with endogenous route, cross-transmission and environmental contamination.** The results are displayed for importation probability *f*, sensitivity parameter *ϕ*, log-likelihood and relative contributions *R*_*i*_ where *i* ∈ {endogenous, cross-transmission, environment}.

**S24 Fig. Pairwise plots of samples from the posterior distribution for the transmission parameters of the full model.** The plots were generated from the data of ICU A before renovation using the full model with endogenous route, cross-transmission and environmental contamination.

**S25 Fig. Coverage probabilities for the submodel using Jeffreys prior.** (a) - (b) ICU A before and after renovation, respectively. (c) - (d) ICU B before and after renovation, respectively.

**S26 Fig. Coverage probabilities for the full model using Jeffreys prior.** (a) - (b) ICU A before and after renovation, respectively. (c) - (d) ICU B before and after renovation, respectively.

